# N-linked surface glycan biosynthesis, composition, inhibition, and function in cnidarian-dinoflagellate symbiosis

**DOI:** 10.1101/820894

**Authors:** Trevor R. Tivey, John Everett Parkinson, Paige E. Mandelare, Donovon A. Adpressa, Wenjing Peng, Xue Dong, Yehia Mechref, Virginia M. Weis, Sandra Loesgen

## Abstract

The success of symbioses between cnidarian hosts (e.g. corals and sea anemones) and micro-algal symbionts hinges on the molecular interactions that govern the establishment and maintenance of intracellular mutualisms. As a fundamental component of innate immunity, glycan-lectin interactions impact the onset of marine endosymbioses, but our understanding of the effects of cell surface glycome composition on symbiosis establishment remains limited. In this study, we examined the canonical N-glycan biosynthesis pathway in the genome of the dinoflagellate symbiont *Breviolum minutum* (family Symbiodiniaceae) and found it to be conserved with the exception of the transferase GlcNAc-TII (MGAT2). Using coupled liquid chromatography-mass spectrometry (LC-MS/MS), we characterized the cell surface N-glycan content of *B. minutum*, providing the first insight into the molecular composition of surface glycans in dinoflagellates. We then used the biosynthesis inhibitors kifunensine and swainsonine to alter the glycan composition of *B. minutum*. Successful high-mannose enrichment via kifunensine treatment resulted in a significant decrease in colonization of the model sea anemone Aiptasia (*Exaiptasia pallida*) by *B. minutum*. Hybrid glycan enrichment via swainsonine treatment, however, could not be confirmed and did not impact colonization. We conclude that functional Golgi processing of N-glycans is critical for maintaining appropriate cell surface glycan composition and for ensuring colonization success by *B. minutum*.

## Introduction

The biodiversity and productivity of coral reef ecosystems relies on the interactions of cnidarian hosts with dinoflagellates in the family Symbiodiniaceae (formerly the genus *Symbiodinium* [1]). These unicellular endosymbionts provide photosynthate to their hosts in return for inorganic carbon and nitrogen and a high light environment which enable the partners to thrive in nutrient-limited waters. In most symbiotic cnidarian species, each generation a host must first engulf potential symbionts, select algae for productive symbiosis in the symbiosome (a modified host vacuole), and allow for proliferation within its tissue. The biological steps that regulate this colonization process are therefore critical to symbiosis establishment and homeostasis. Innate immunity pathways govern the onset of symbiosis when algae first come into contact with their future host cells [2, 3]. Cnidarians rely on expanded innate immune repertoires for defense against pathogens and selection of microbes [4–8]. These immune pathways are often co-opted targets for microorganisms looking to benefit from a host, and therefore offer potential avenues for symbiont coevolution and entry into hosts [9].

The surfaces of most eukaryotic cells are decorated with a complex array of surface sugars that play a major role in cell-cell communication [10]. These cell surface glycans are recognized by carbohydrate-binding proteins, known as lectins, to mediate events such as cell homing, sperm-egg binding, and host-microbe interactions [11–13] (Figure 1). Glycans are also a major category of microbe-associated molecular pattern that interact with host pattern recognition receptors that, in turn, regulate innate immunity and symbiosis [2, 14]. Lectins on host cells have the ability to either promote microbial attachment and colonization or to block the interactions required for entry into the host [15–18]. These glycan-lectin interactions form a critical component of host innate immunity that functions in self/non-self discrimination and pathway activation against potential pathogens [19, 20].

**Figure 1.**
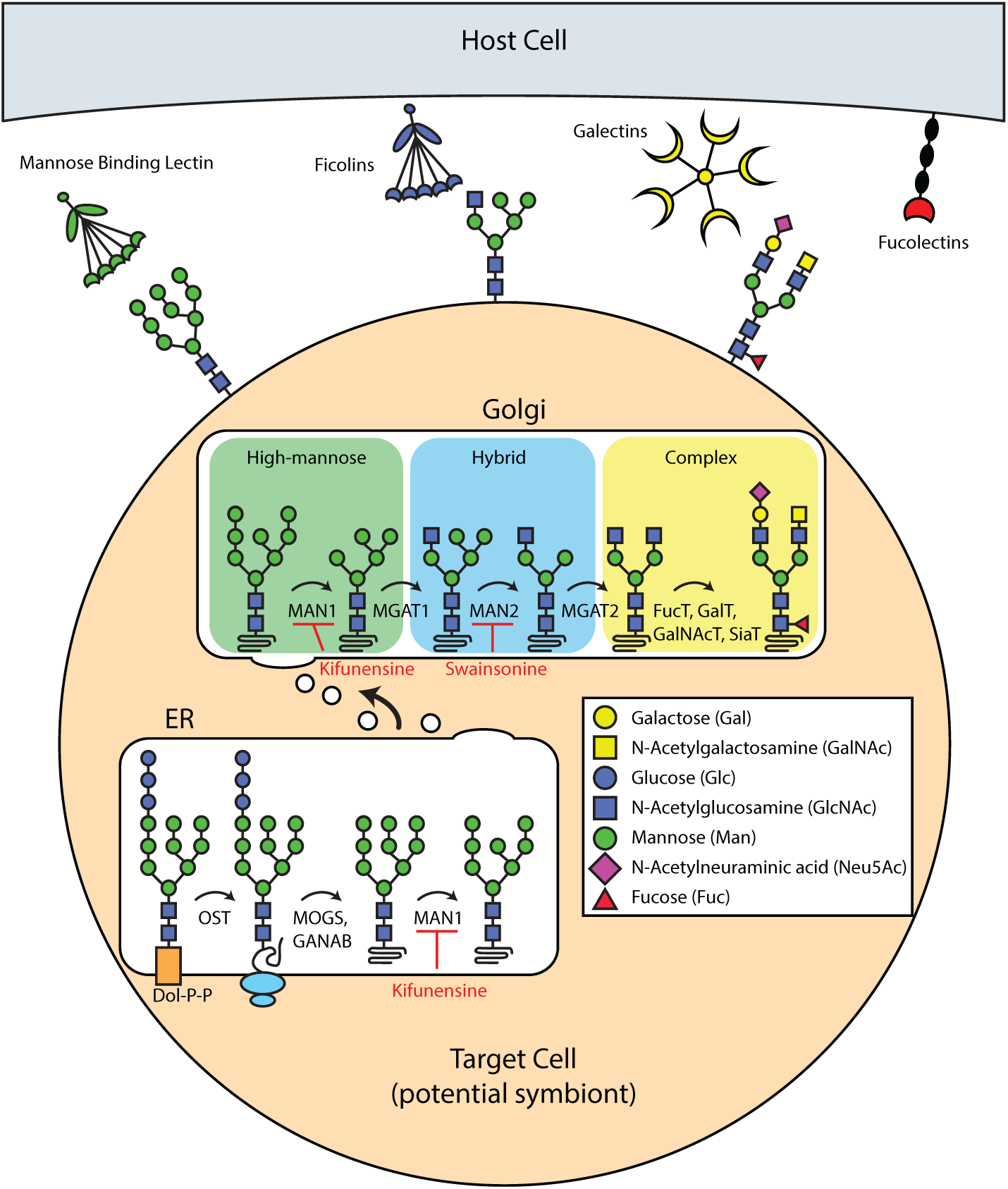
Canonical N-glycan biosynthesis and N-glycan recognition in animals. The biosynthesis of N-glycans begins with the transfer of a nucleoside activated N-acetylglucosamine (GlcNAc) to a dolichol membrane anchor (Dol-P-P). A fourteen-sugar precursor structure is created by elongation, in the ER lumen. This structure is then mass-transferred to an Asn residue of a protein undergoing synthesis and aids in protein folding. N-glycans are further processed by glucosidases (MOGS, GANAB) to remove glucose, and then by ER mannosidase I (MAN1) to remove mannose residues. MAN1 enzymes act in both the ER and the Golgi to transform oligomannoside into hybrid glycans by N-acetylglucosaminyltransferase 1 (MGAT1). To create complex glycans, glycoproteins in the Golgi are further cleaved by mannosidase II (MAN2), then a series of transferases add GlcNAc, galactose, and other sugar residues to create complex N-glycans. Small molecules that act as enzymatic inhibitors can be used to block parts of this pathway, for example kifunensine and swainsonine. Pattern recognition receptors (PRRs) from host cells (Mannose-binding lectin, ficolins, etc.) interact with surface glycans on target cells to mediate immunity. In dinoflagellate symbionts, we have limited knowledge regarding the conservation of this general N-glycan biosynthesis pathway found in animals and plants. This N-glycan biosynthesis pathway was adapted from Helenius and Aebi [82].

While the importance of glycan recognition has been well documented in some host-symbiont associations, the molecular characterization of surface glycans in the family Symbiodiniaceae and their function in symbiosis is just beginning to be uncovered. Other alveolates offer potential insight into glycan-lectin interactions in Symbiodiniaceae mutualisms [21]. For example, the well-studied apicomplexan parasites *Plasmodium falciparum* and *Toxoplasma gondii* have simplified glycomes that lack the longer immunogenic glycans often detected by host immunity [22–24]. The alveolate parasite *Perkinsus marinus*, a member of the sister phylum to dinoflagellates, has glycans that are recognized during shellfish infection by a host galectin receptor [25]. The prevalent nature of glycan-lectin interactions in parasitic groups that are closely related to dinoflagellates suggests that the glycomes of Symbiodiniaceae may also have evolved to gain access to the host cnidarian environment.

The strongest evidence supporting glycan-lectin recognition during the onset of Symbiodiniaceae mutualisms comes from experimental manipulations of symbiont cells through enzymatic cleavage of surface glycans. Several species of Symbiodiniaceae have been experimentally de-glycosylated, leading to reduced symbiont uptake by larval, juvenile, and adult aposymbiotic animals [26–29]. Though the effects are not entirely consistent from study to study, the data suggest that N-glycans in particular are an important determinant of how rapidly hosts are colonized by symbionts. It remains uncertain whether disruption in the onset of symbiosis is driven primarily by changes in the overall abundance of N-glycans (i.e. quantity) or by changes in the composition of N-glycans (i.e. complexity), particularly the relative proportions of high-mannose, hybrid, and complex N-glycans.

Although recent advances in the field of glycomics have made the direct chemical analysis of carbohydrates feasible, most studies probing the Symbiodiniaceae cell surface have relied on indirect lectin binding to characterize glycan type and abundance, and to mask glycan interactions. For example, a variety of fluorescent FITC-labeled lectin probes were used to examine the cell surface of *Cladocopium* sp. (type C1f) isolated from the coral *Fungia scutaria* [29]. The C1f symbionts were strongly labeled with Concanavalin A (ConA) and Jacalin lectins, which bind *α*-mannosyl, *α*-glucosyl and *α*-galactosyl (Jacalin only) groups. Logan et al. (2010) further examined other genera of Symbiodiniaceae, finding that binding varied by species. *Breviolum minutum* in particular was labeled by a diversity of lectins recognizing mannose, glucose, galactose, N-acetylglucosamine (GlcNAc), and N-acetylgalactosamine (GalNAc) residues. In contrast, Parkinson et al. (2018) used a glycan-binding array of 40 lectins and found only small differences in the glycan profiles between the free-living species *Symbiodinium pilosum* and the mutualistic species *B. minutum*. Both studies provide evidence that Symbiodiniaceae produce surface glycoproteins containing high-mannose as well as hybrid/complex type N-glycans, but it remains unclear which types of N-glycans specifically contribute to recognition for colonization by mutualistic species. Here, we employed liquid chromatography coupled mass spectrometry (LC-MS/MS) to characterize *B. minutum*’s cell surface N-glycan content on the molecular level.

The N-glycan biosynthesis pathway is well-described in yeast and humans, and a number of natural products have been isolated that serve as inhibitors [30, 31]. If the canonical N-glycan pathway found in animals and plants is conserved in dinoflagellates (as shown in Figure 1), it should be possible to use commercially-available inhibitors to alter glycan biosynthesis in Symbiodiniaceae by primarily changing the composition of the glycome instead of altering the overall abundance of N-glycans. Kifunensine is one such molecule, a plant-derived alkaloid, that inhibits the activity of ER Mannosidase I [32]. Because blocking cleavage prevents the synthesis of hybrid and complex glycan structures, this inhibition results in an increase in the proportion of high-mannose type N-glycans (Figure 1) [33, 34]. Swainsonine is another small molecule inhibitor that blocks later stage activity of Golgi Mannosidase II [35]. Therefore, swainsonine treatment increases the proportion of hybrid-type N-glycan content and decreases the proportion of complex-type glycans (Figure 1) [36, 37]. These two inhibitors have the potential to differentially modify symbiont surface glycan composition allowing one to experimentally test for preferential uptake by cnidarian hosts of algal cells with certain surface glycan profiles.

In this study, we directly determined the structure and abundance of N-glycans from symbiotic *B. minutum*, used small molecule inhibitors to manipulate the glycome of *B. minutum*, and subsequently tested colonization success of these altered algae in model cnidarian hosts.

## Materials and methods

Raw data and R code for the following analyses can be found in Supplementary File S1. For complete details regarding colonization and image analysis, protocols can be accessed on protocols.io at dx.doi.org/10.17504/protocols.io.j8ucrww.

### Identification of N-Glycan Biosynthesis Homologs

Our hypotheses were based on the idea that Symbiodiniaceae share the canonical N-linked glycosylation biosynthesis pathway with other eukaryotes. To strengthen this premise, we first searched for conservation of dolichol-linked precursor oligosaccharide synthesis and ER/Golgi processing pathways in Symbiodiniaceae genomes. We used *Homo sapiens* protein sequences to search for predicted proteins in the Symbiodiniaceae genomes of *Breviolum minutum* [38], *Symbiodinium microadriaticum* [39], and *Fugacium kawagutii* [40]. The genomes of *Perkinsus marinus* strain ATCC 50983 [41], *Plasmodium falciparum* strain 3D7 [42], and *Toxoplasma gondii* strain ME49 [43] were also included as closely-related alveolate comparisons. In total, 34 manually annotated protein sequences from UniProtKB were used as queries representing proteins involved in N-glycan precursor synthesis and N-glycan processing in the ER and Golgi [44]. Enzymes that have similar functions but that are involved in degradation were included as well, such as lysosomal *α*-mannosidase and *β*-hexosaminidase 1. Predicted proteins for Symbiodiniaceae species were searched on reefgenomics.org, a platform hosting the genomes of several marine organisms including three Symbiodiniaceae genomes [45]. Predicted proteins for the three other alveolate genomes were searched on protists.ensembl.org, a browser for protist genomes [46]. Homologous proteins were considered to be present in a genome if the top protein BLAST hit exceeded the following cutoff: eval < 1 x 10^-5^, bitscore > 50, identity score > 25%. T-Coffee was used to align amino acid sequences of putative *α*-mannosidases from Symbiodiniaceae against *α*-mannosidase 2 and lysosomal *α*-mannosidase [47, 48]. Sequences were chosen from *Drosophila melanogaster*, *Homo sapiens*, *Mus musculus*, and *Arabidopsis thaliana* to allow for comparison to previous studies characterizing the active sites of *α*-mannosidase [37, 49].

### Treatment of Algal Cultures

*Breviolum minutum* (strain Mf1.05b), a homologous or native symbiont found in Aiptasia, was used for all experiments. Algal cultures were maintained in f/2 media in an incubator at 26°C under a 12:12 light:dark cycle and a light intensity of 50 µmol quanta • m^-2^ • s^-2^. For experimental treatment, algal cultures were reconstituted at 1 x 10^6^ cells • mL^-1^ in f/2 media. To cleave N-glycans for dosage and colonization experiments, algal cultures were incubated in a range of concentrations of Peptide:N-glycosidase F for 24 h prior to flow analysis (500, 100, 50, 5, and 0 U PNGase F • mL^-1^ f/2 media). For biosynthesis inhibition treatment, kifunensine and swainsonine stocks were prepared at concentrations of 50 mM in 100% dimethyl sulfoxide (DMSO) solvent. Cultures were incubated for 1 wk in 1 mM kifunensine or swainsonine in 2% DMSO in f/2 (a 1:50 dilution) or in a 2% DMSO in f/2 vehicle control which was well-tolerated. Pilot experiments tested a range of inhibitor concentrations from 10 µM to 10 mM and tested a range of DMSO concentrations from 2-10% (data not shown.)

### Identification of N-Glycans via ESI LC-MS/MS

*Breviolum minutum* (strain Mf1.05b) was cultured as described above. For enzymatic treatment, 1 L of culture was spun down and treated with Peptide:N-glycosidase F for 72 h (50 U PNGase F mL^-1^ f/2 media). The reactions were stirred at 37 C for 72 h. The glycan sample was dialyzed using 10k MWCO dialysis tubing in 100 mL ultrapure water; water was changed three times over the course of 48 h. The glycan sample was lyophilized and 1 mg of lyophilized sample was reduced and permethylated as previously reported [50–53]. In summary, lyophilized samples were dissolved in 10 µL of borane-ammonia complex solution (10 mg• mL^-1^) and incubated in the water bath at 60°C for 1h. After incubation, 1 mL of methanol was added to the samples and vacuum dried (LABCONCO Cor. Kansas City, MO). A 1 mL aliquot of methanol was repeatedly added and dried until borate was completely removed by co-evaporating with methanol. The reduced sample was redissolved in a 30 µL aliquot of dimethyl sulfoxide (DMSO), 1.2 µL water, and 20 µL iodomethane. Sodium hydroxide beads (Sigma, St. Louis, MO; suspended with DMSO) were packed into a spin column, centrifuged at 1800 rpm for 2 min, and then washed with 200 µL of DMSO. Samples were loaded onto the column and incubated at room temperature for 25 min. Another 20 µL aliquot of iodomethane was added into the column and incubated for 15 min. After incubation, permethylated glycans were collected by centrifuging at 1800 rpm for 2 min. Next, 30 µL of acetonitrile (ACN) was added into the column and centrifuged again to collect the eluent. The ACN eluent was combined with previous DMSO solution and dried completely.

Reduced and permethylated glycans were then subject to nano-C18-LC-MS/MS analysis. LC-MS analysis was performed on a Dionex 3000 Ultimate nano-LC system (Dionex, Sunnyvale, CA) interfaced to an LTQ Orbitrap Velos mass spectrometer (Thermo Scientific, San Jose, CA). A 200 ng aliquot of the lyophilized glycans were injected. Glycans were first retained on a trap column (Acclaim PepMap 100, 75 μm × 2 mm, 3 μm, Thermo Scientific) and then separated by a nanoC18 column (Acclaim PepMap 100, 75 μm × 150 mm, 2 μm, Thermo Scientific). The flow rate was 0.35 µL• min^-1^. The gradient was attained using mobile phase A (98% water, 2% ACN, 0.1% formic acid) and mobile phase B (ACN, 0.1% formic acid) employing the following multi-steps: 20% B for 10 min; 20% - 42% B from 10 to 11 min; 42% - 55% B from 11 to 48 min; 55% - 90% B from 48 to 49 min; 90% B from 49 to 54 min; 90% - 20% B from 54 to 55 min; 20% B from 55 to 60 min.

The resolution of LTQ Orbitrap Velos was set to 100,000. Collision induced dissociation (CID) was applied to achieve tandem mass spectrometry data (MS^2^) with an injection time of 10 ms, collision energy of 35%, activation Q of 0.25. The data dependent acquisition (DDA) method was used with the top 4 ions detacted in the full MS to be picked up for CID fragmentation. The identification and quantitation of glycans were performed by Multiglycan software [54] and then manually checked.

### Lectin Labeling and Flow Cytometry

After treatment, 100 µL algal samples were taken from each treatment, rinsed in 3 mL of filtered artificial seawater (FSW) and centrifuged in 75 mm borosilicate glass tubes (1670 rpm for 5 min). Pellets were resuspended in 50 µL of 3.3X PBS, prepared from a 10X PBS stock solution (0.02 M NaH_2_PO_4_, 0.077 M Na_2_PO_4_, 1.4 M NaCl, pH 7.4). Subsequently, 50 µL of 2X labeling solution containing lectins conjugated to phycoerythrin (PE) fluorophores was added to each sample at a final concentration of 5 µg lectin • mL^-1^ PBS. Lectins were coupled to PE using the Lightning-Link R-PE Antibody Labeling Kit (Novus Biologicals #703-0010); PE was chosen in order to avoid any overlap with algal autofluorescence. Cyanovirin-N lectin (CVN-PE) was used to label high-mannose N-glycans. *Microcystis viridis* lectin (MVL-PE) was used to label core N-glycan residues composed of GlcNAc and mannose [55]. *Phaseolus vulgaris* Phytohaemagglutinin lectin (PHA-L-PE) was used to label complex glycans. Their respective glycan recognition profiles summarized from publicly available glycan array data are shown in Supplementary Figure 1 (http://www.functionalglycomics.org). These lectins were chosen for their micromolar binding affinities and comparatively greater selectivity and cleaner binding profiles. The surface glycan composition of labeled algal symbionts was measured on a CytoFLEX 5L flow cytometer. Algal populations were identified using forward and side scatter, and doublets were removed using FSC-Width in order to avoid overestimating fluorescent intensities of cells. A minimum of 20,000 cells labeled with PE-conjugated lectins were excited at 561 nm and emission was captured with a 585/42 nm bandpass filter. The median fluorescence intensity for each FSC-Width population was averaged for each treatment and compared using ANOVA. The arithmetic mean and geometric mean fluorescence intensity were also averaged by treatment and analyzed.

### Colonization Experiments

Aposymbiotic adult Aiptasia polyps were obtained after repeated menthol bleaching [56] and maintenance in the dark. Prior to the beginning of experiments, polyps were individually plated in 24-well plates and exposed to the same 12:12 light:dark cycle as algal cultures for 1 wk. Animals were then visually checked under an inverted compound fluorescence microscope for the presence of any contaminating symbionts, and only those completely lacking symbionts were used in colonization experiments. Algae from each treatment were rinsed in FSW and resuspended to 2 x 10^6^ cells • mL^-1^. Each anemone was inoculated with 1 x 10^6^ cells (0.5 mL of algal treatment) in a total volume of 1 mL FSW. After 48 h, the wells were rinsed three times with FSW, and left for 24 h before overnight fixation in 4% paraformaldehyde in 1X PBS. To prepare for imaging symbiont density, the oral discs with tentacles were dissected from the columns of each polyp and mounted on slides in a mounting solution (90% glycerol, 10% 1X PBS) as described previously [57].

### Image Analysis

To normalize symbiont density to host surface area, epifluorescence microscopy was used to capture autofluorescence from the symbionts and the host anemone tissue as detailed previously [57]. Symbiont chlorophyll autofluorescence was captured under the Cy3 (red) filter, whereas autofluorescence from host green fluorescence was captured using the GFP (green) filter. Images of symbiont density after colonization of hosts were captured with a Zeiss AxioObserver A1 microscope and an Axiovert ICm1 camera (Carl Zeiss AG, Jena, Germany). Algal cell numbers were automatically quantified by setting a predetermined cell radius for the ITCN plugin for Fiji (ImageJ2). To calculate algal density as a proxy for symbiont colonization, the total algal cell number was divided by the surface area of host autofluorescence.

## Results and Discussion

### N-Glycan Biosynthesis Pathway Conservation in Alveolates

We hypothesized that the pathways underlying N-linked glycoprotein production in Symbiodiniaceae are similar to those in humans and other well-studied eukaryotes. Our findings support this hypothesis. We found a large degree of conservation among members of these groups using BLASTP searches for homologous proteins. We found strong evidence for N-glycan biosynthesis conservation up to MGAT1, an N-acetylglucosaminyltransferase essential for hybrid and complex type glycan processing (Table 1, Figure 1). Golgi *α*-mannosidase 2, the next step in the N-glycan pathway, also appears in a BLASTP search of Symbiodiniaceae genomes, however the sequence hits were more similar to lysosomal *α*-mannosidase, especially in conserved regions (Supplementary Figure 2, Supplementary Table 1). Without functional or subcellular localization data, it is difficult to discern whether this *α*-mannosidase functions during N-glycan biosynthesis or only during degradation; regardless, it should be inhibited by swainsonine based on conserved residues (Supplementary Table 1) [37]. Although all Symbiodiniaceae appear to be missing GlcNAc-TII (MGAT2), the *B. minutum* genome includes transferases GlcNAc-TIII (MGAT 3), GlcNAc-TIV (MGAT 4) and GlcNAc-TV (MGAT5), which are required for bisecting, tri-, and tetra-antennary N-glycans (Table 1). The presence of these transferases supports the existence of hybrid glycan biosynthesis in Symbiodiniaceae. With the absence of MGAT2, it remains unclear whether the *α*-1-6 branch required for complex glycans can be formed [31]. In addition to the majority of enzymes required for N-glycan branching, Symbiodiniaceae genomes contain N-glycan transferases for xylose, fucose, and galactose (Table 1, Supplementary Table 2). These moieties are possible targets for the expanded set of cnidarian ficolin-like proteins, fucolectins, and c-type lectins found in Aiptasia [58, 59]. Nevertheless, these Symbiodiniaceae species appear to have the capacity to create high-mannose, hybrid, and potentially complex glycans.

**Table 1:** N-glycan biosynthesis in alveolates.

| **Description** | **Protein** | **B.**<br>**min.** | **S.**<br>**mic.** | **F.**<br>**kaw.** | **P.**<br>**mar.** | **T.**<br>**gon.** | **P.**<br>**fal.** |
| --- | --- | --- | --- | --- | --- | --- | --- |
| **Dol-P precursor synthesis** |  |  |  |  |  |  |  |
| UDP-N-acetylglucosaminyltransferase subunit | ALG13 | + | + | + |  | + | + |
| UDP-N-acetylglucosaminyltransferase subunit | ALG14 | + | + |  | + | + | + |
| Mannosyltransferase-1 | ALG1 | + | + |  | + | + |  |
| α-1,3-mannosyltransferase | ALG2 | + | + | + | + | + | |
| α-1,2-mannosyltransferase | ALG11 | + | + | | + | + | |
| α-1,3-mannosyltransferase | ALG3 | + | + | | + | | |
| α-1,2-mannosyltransferase | ALG9 | + | + | + | + | | |
| α-1,6-mannosyltransferase | ALG12 | | + | + | + | | |
| DolP-glucosyltransferase | ALG5 | + | + |  | + | + |  |
| α-1,3-glucosyltransferase | ALG6 | + | + | + | + | + | |
| α-1,3-glucosyltransferase | ALG8 | + | + | + | + | + | |
| α-1,2-glucosyltransferase | ALG10 | + | + | + | | + | |
| Oligosaccharyl transferase subunit STT3 | STT3 | + | + | + | + | + | + |
| **ER Processing** |  |  |  |  |  |  |  |
| Mannosyl-oligosaccharide glucosidase | GCS1 | + | + |  | + | + |  |
| Neutral α-glucosidase AB | GANAB | + | + | + | + | + | |
| ER α-1,2-mannosidase | MAN1B1 | + | + | + | + | | |
| **Golgi Processing** |  |  |  |  |  |  |  |
| α-1,2-mannosidase IA | MAN1A1 | + | + | + | + | | |
| α-1,2-mannosidase IB | MAN1A2 | + | + | + | + | | |
| (α-1,3) 2-β-GlcNAc transferase I | MGAT1 | + | + | + | + | | |
| α-mannosidase 2 | MAN2A1 | | | | | | |
| Lysosomal α-mannosidase | MAN2B1 | + | + | + | + | | |
| (α-1,6) β-1,2-GlcNAc transferase II | MGAT2 | | | | | | |
| (β-1,4) β-1,4-GlcNAc transferase III | MGAT3 | + | | | + | | |
| (α-1,3) β-1,4-GlcNAc transferase IVC | MGAT4C | + | + | | | | |
| (α-1,6) β-1,6-GlcNAc transferase V | MGAT5 | + | + | | | | |
| **Complex** |  |  |  |  |  |  |  |
| Xylosyltransferase 1 | XYLT1 | + | + |  |  |  |  |
| α-1,6-fucosyltransferase 8 | FUT8 | | + | + | | | |
| α-1,3-fucosyltransferase 11 | FUT11 | + | + | | + | | |
| α-1,4-fucosyltransferase 13 | FUT13 | + | + | | | | |
| β-1,4-galactosyltransferase 1 | B4GALT1 | + | + | + | | | |
| β-1,3-galactosyltransferase 1 | B3GALT1 | + | + | + | + | | |
| β-galactoside α-2,6-sialyltransferase 1 | SIAT1 | | | | | | |
| β-galactoside α-2,6-sialyltransferase 2 | SIAT2 | | | | | | |
| **Paucimannose** |  |  |  |  |  |  |  |
| β-hexosaminidase 1 | HEXO1 | + | + | + | + | | |

The N-glycan biosynthesis pathway BLASTP protein hits for the Symbiodiniaceae species (*Breviolum minutum*, *Symbiodinium microadriaticum*, and *Fugacium kawagutii*) were most similar to the closely related *Perkinsus marinus*, an endoparasite of oysters (Table 1). Among the Symbiodiniaceae, we found the N-glycan biosynthesis pathway in *F. kawagutii* to be the least conserved, which agreed with the initial, less stringent analysis of the N-glycan pathway from the *F. kawagutii* draft genome [40]. This lack of conservation could reflect its phylogenetic position (one of the most derived lineages within the group) and/or ecology: of the three species whose genomes were searched, it is the only one that is exclusively free-living and nonsymbiotic. If hybrid and/or complex N-glycan production is a key to symbiosis establishment and maintenance, there may be less selective pressure to retain these genes in nonsymbiotic species. However, as the Symbiodiniaceae genomes are still in draft form, the absence of pathways may be a result of differences in sequencing and/or annotation completeness, especially considering the decreased number and shorter length of gene models in the original *F. kawagutii* genome [40, 60]. In examining more evolutionarily distant comparisons, *B. minutum* and *S. microadriaticum* appear to have a greater capacity for N-glycan biosynthesis than many other algal protists, including other dinoflagellates, chlorophytes, diatoms, and haptophytes [40].

The degree of conservation in the N-glycan biosynthesis pathway of the Symbiodiniaceae is striking when compared to the degenerate pathways of the alveolate parasites *Plasmodium falciparum* and *Toxoplasma gondii*. In contrast to the Symbiodiniaceae, *P. falciparum* has severely limited N-glycosylation and O-glycosylation, and is only able to produce the chitobiose core using ALG13 and ALG14 (Table 1) [22, 61]. Instead, *P. falciparum* relies on GPI anchors (glycolipids) to stabilize proteins, and uses its own lectins to bind to host cells [62]. The *T. gondii* glycome is predominantly composed of high-mannose (Man_5–8_(GlcNAc)_2_) and shorter paucimannose (Man_3–4_(GlcNAc)_2_) glycans [23]. These degenerate N-glycans are partially a consequence of a loss of ALG glycosyltransferases in both parasites [24] (Table 1). The general conservation of precursor synthesis and ER/Golgi processing in the Symbiodiniaceae and *P. marinus* suggests strong differences in N-glycan processing between apicomplexans and Dinozoans (containing dinoflagellates and their sister phylum Perkinsozoa), supporting the hypothesis that among alveolates, only apicomplexans have undergone secondary loss of ALG enzymes [22].

### N-Glycan Identification in *Breviolum minutum*

With the knowledge that N-glycan biosynthesis pathways are present and even conserved in *Breviolum minutum*, we sought to characterize the N-linked glycome. We harvested algal cells, cleaved the N-linked surface glycans, and after per-methylation used LC-MS/MS for glycan analysis [50–53, 63]. Based on molecular weight, retention time, and comparison to carbohydrate standards, the overall cleaved N-linked glycan sample exhibited 52% high-mannose glycans, 12% core-fucosylated glycans, and, to our surprise, 3% sialylated glycans (Figure 2a). Overall, 29 glycan structure were identified in the liquid-chromatography coupled mass spectrometry analysis (Figure 2b). While the enrichment of high-mannoside glycans has been shown indirectly in *B. minutum* before, this is the first direct, analytical verification of the result [28, 29, 57, 64].

**Figure 2.**
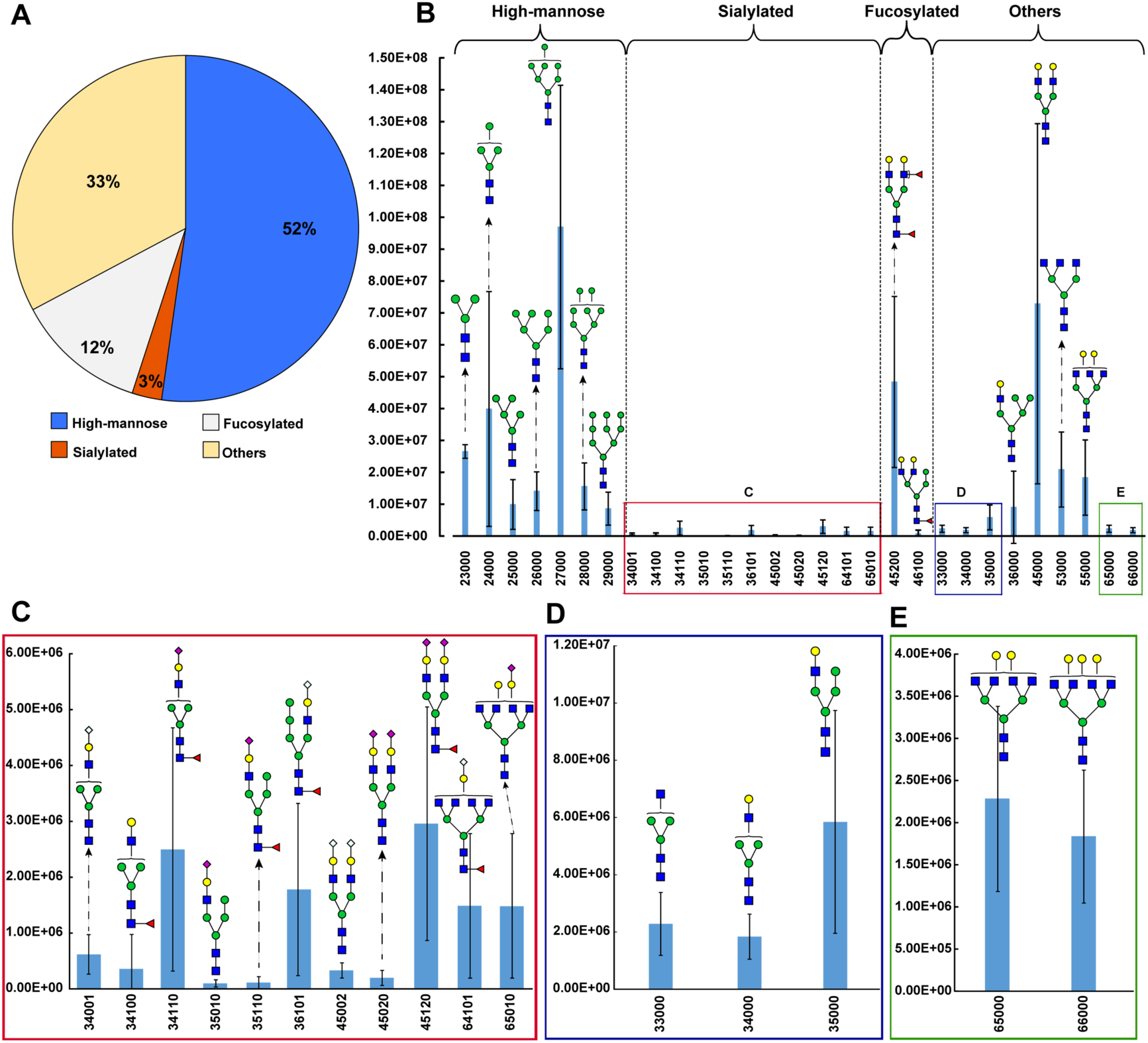
Quantitative glycomic analysis in *Breviolum minutum* via LC-MS/MS. **(a)** Proportions of different N-glycan types represented in a sample of enzymatically cleaved *B. minutum* surface glycans. **(b)** Absolute quantitation of N-glycans and their putative structures. Boxed sections indicate zoomed in areas with 20x **(c)** and 10x magnification **(d, e)**. Symbol: blue square – N-Acetylglucosamine (GlcNAc); green circle –mannose (Man); yellow circle – galactose (Gal); purple diamond – N-acetylneuraminic (sialyic) acid (NeuAc); hollow diamond – N-Glycolylneuraminic acid (NeuGC); red triangle – fucose (Fuc). Nomenclature: a five digit code is used to represent glycan structures. The 1^st^ number represents the number of HexNAc (GlcNAc); the 2^nd^ number represents the number of Hex (Man + Gal); the 3^rd^ number represents the number of DeoxyHex (Fuc); the 4^th^ number represents the number of sialic acid (NeuAc); the 5^th^ number represents the number of N-Glycolylneuraminic acid (NeuGC). For example, 45120 represents HexNAc_4_Hex_5_DeoxyHex_1_NeuAc_2_.

Sialic acid (also called N-acetylneuramic acid) is often considered a hallmark of the human glycome as it can be found mostly on cell surfaces of ‘higher’ invertebrates. In humans, sialic acid modulates a variety of normal and pathological processes. Many pathogens bind to it (e.g. human *Influenza* A, *Vibrio cholerae*, *Plasmodium falciparum*, *Heliobacter pylori*) or express it on their surfaces to interact with human cells (Neisseria spp., *Trypanosoma cruzi*) [65]. Sialic acid is rarely found in algae and this is the first report of sialic acid in dinoflagellates [66–68]. Future studies should test whether sialic acid is a crucial component of cnidarian-dinoflagellate symbiosis establishment.

### Effects of N-Glycan Removal on Colonization Success in Host Anemones

We expected that glycosidase treatment with PNGase F would decrease the total number of N-linked glycans on the cell surface of *B. minutum* and subsequently reduce colonization success. We measured changes to the algal cell surfaces using fluorescently tagged lectin (CVN-PE) that recognizes glycans highly enriched with *α*(1-2)-linked dimannosides. After exposure to PNGase F, CVN-PE labeling mean fluorescence intensities decreased, indicating that fewer high-mannoside glycans were present on algal cell surfaces after treatment (Figure 3a; ANOVA, Tukey post-hoc test, *p* < 0.001). Aiptasia polyps were then inoculated with glycosidase-treated or untreated cells. After initial colonization, symbiont density was lower in hosts exposed to the cells with reduced N-glycan abundances (Figure 3b; t-test, *p* < 0.05).

**Figure 3.**
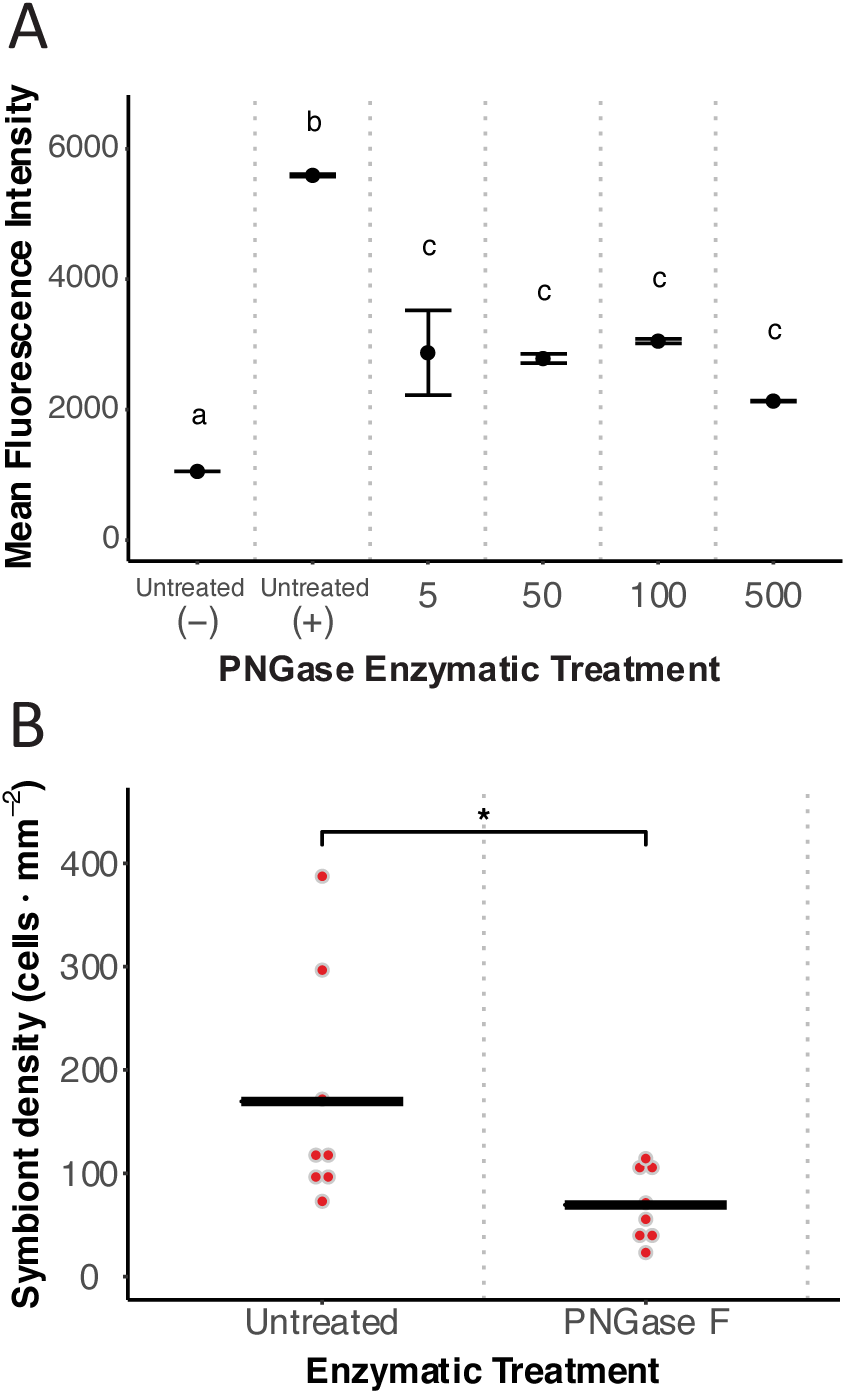
Effect of PNGase F treatment on *Breviolum minutum*. **(a)** Cell surface N-glycan removal. After 48 h of PNGase F treatment at different concentrations, glycan abundance on cell surfaces was measured through the binding of fluorescently tagged lectin CVN. Unique letters represent significant differences in mean fluorescence intensity following ANOVA (*p* < 0.001; n = 2 replicates per treatment). Untreated cells without fluorescently-labeled lectins were used as a negative control (-), while untreated cells with fluorescently-labeled lectins were used as a positive control (+). Error bars indicate total range. **(b)** Colonization rates of untreated and PNGase F-treated algae in adult polyps. Dots represent data from replicate polyps. Asterisks indicate a significant difference between treatments (*t*-test, *p* < 0.05).

These results support the importance of N-glycans in host-symbiont interactions during the onset of symbiosis and are consistent with previous studies in Aiptasia and coral larvae. N-glycosidase treatments of algal symbionts blocked colonization of Aiptasia adults by algae, as did treatments with O-glycosidase, trypsin, and *α*-amylase [28]. Similarly, N-glycosidase and trypsin treatments inhibited symbiont colonization in larvae of the coral *Fungia scutaria* [29]. In contrast, *α*-amylase had no effect on the percentage of *F. scutaria* larvae colonized or in algal density within individual larvae. In another coral larval study using *Acropora tenuis*, *α*-amylase and trypsin increased the number of symbiont cells taken up by larvae [26]. Across all enzymatic treatments to date, N-glycosidases have had the most consistent negative effect on colonization rates of Symbiodiniaceae in cnidarian hosts. Enzymatic treatment with trypsin and PNGase F has also been used to decrease affinity of isolated cnidarian lectins to their glycan binding sites on Symbiodiniaceae, supporting the hypothesized function of glycan-lectin interactions in symbiosis [69, 70]. This consistency across many studies suggests that the overall abundance of N-glycans on the Symbiodiniaceae cell surface impacts the rate of symbiont uptake by cnidarian hosts.

### N-Glycan Biosynthesis Manipulation in *Breviolum minutum*

After confirming the effect of glycan abundance on colonization success, we were interested in understanding how the composition of different types of N-glycans affected the onset of symbiosis. To alter glycan composition, we inhibited the N-glycan biosynthesis pathway at ER and Golgi mannosidase enzymatic steps using the small molecule inhibitors kifunensine and swainsonine (Figure 4a). To confirm that the manipulations altered the composition rather than the abundance of N-glycans, we labeled cells with MVL-PE, which binds to structures consisting of Manβ(1-4)GlcNAcβ(1-4), a common core found in all three N-glycan types (Figure 4b). MVL-PE labeling showed a small but statistically significant increase in median fluorescence intensity in the kifunensine treatment compared to untreated cells (Figure 4c; ANOVA, Tukey post-hoc test, *p* < 0.001), but showed no difference between arithmetic mean (ANOVA, Tukey post-hoc test, *p* = 0.99) and geometric mean (ANOVA, Tukey post-hoc test, *p* = 0.65) fluorescence intensities. In general there were similar N-glycan abundances between all treatments. This relative stability in glycan abundance allowed us to analyze the remaining lectin binding data as primarily representative of differences in glycan composition.

**Figure 4.**
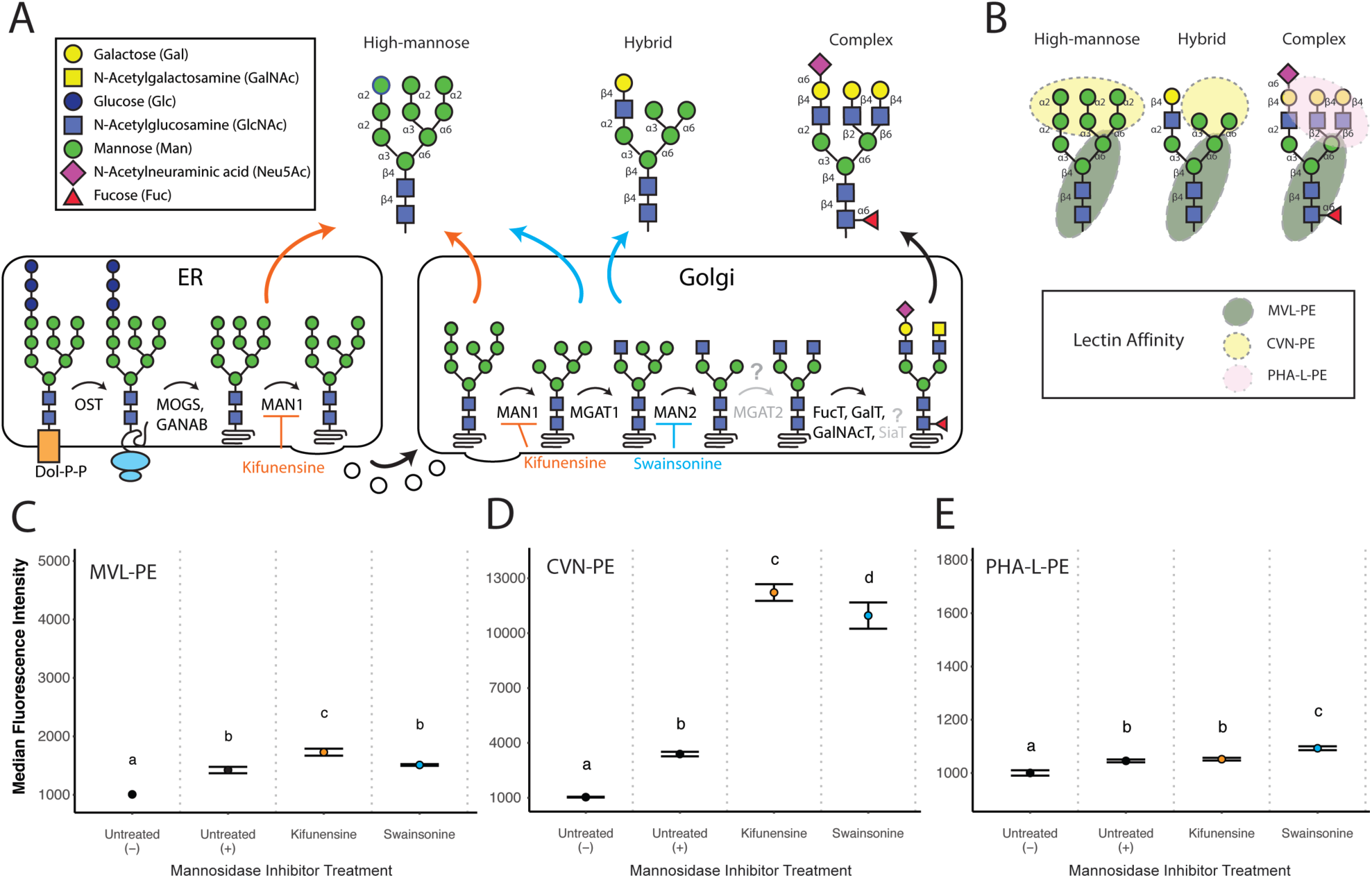
Inhibition of N-glycan biosynthesis in *Breviolum minutum*. **(a)** Pharmacological inhibition of the N-glycan biosynthesis pathway with kifunensine (orange) and swainsonine (teal) in the ER and Golgi and the hypothesized effect on cell surface N-glycans in Symbiodiniaceae cells. Parts of the pathway that lack conservation are shown in gray, indicated by a question mark. **(b)** Glycan recognition sites of the respective lectins within the three types of N-glycans (high-mannose, hybrid, and complex). MVL-PE was used to confirm the core of N-glycans, whereas CVN-PE and PHA-L-PE were used to determine the high-mannose and complex glycan composition, respectively. **(c)** MVL-PE labeling of core N-glycans. Algae that were incubated for one week with kifunensine, swainsonine, or DMSO (control) were labeled with MVL-PE to quantify N-glycan abundance using median fluorescence intensity (n = 3). No differences in mean fluorescence intensity were found between lectin-labeled treatments. **(d)** CVN-PE labeling of high-mannose N-glycans. Treated algae were labeled with CVN-PE to quantify relative amount of high-mannose N-glycans using median fluorescence intensity (n = 3-4). Mean fluorescence intensity comparisons resulted in the same significance. **(e)** PHA-L-PE labeling of complex N-glycans. Treated algae were labeled with PHA-L-PE to quantify relative amount of complex N-glycans using median fluorescence intensity (n = 6-8). Mean fluorescence intensity comparisons resulted in the same significance. Untreated cells without fluorescently-labeled lectins were used as a negative control (-), while untreated cells with fluorescently-labeled lectins were used as a positive control (+). Unique letters represent significant differences in staining percentages (ANOVA, *p* < 0.05).

Kifunensine blocks MAN1, the enzyme responsible for converting high-mannose glycans to hybrid glycans featuring less mannose. To confirm that kifunensine treatment increased the proportion of high-mannose glycans, symbiont cells were labeled with CVN-PE, which binds with α(1-2)-linked dimannosides residing at the branch tips of mannose-rich glycans (Figure 4a). Cells treated with kifunensine had the highest median fluorescence intensity of CVN-PE labeling as expected, significantly greater than both swainsonine-treated and untreated cells (Figure 4d; ANOVA, *p* < 0.05, *p <* 1 x 10^-7^). Swainsonine-treated cells also exhibited significantly greater labeling by CVN for mannosides than untreated cells (Figure 4d; ANOVA, *p <* 1 x 10^-7^). This was expected, as hybrid glycan structures induced by swainsonine should have more abundant mannose moieties than untreated cells that feature a greater proportion of complex, low-mannose glycans (Figure 4a). Analysis of geometric mean fluorescence intensity showed the same significant differences (ANOVA, *p <* 1 x 10^-7^) between every comparison except swainsonine and kifunensine treatments (ANOVA, *p* = 0.28), whereas arithmetic mean fluorescence intensity found no differences between swainsonine and kifunensine treatments (ANOVA, *p* = 0.92) or between swainsonine and untreated cells (ANOVA, *p* = 0.10). Overall, kifunensine had a strong effect on increasing the proportion of high-mannose N-glycans, whereas swainsonine had a weaker effect. The increase in lectin-binding in both treatments supports the presence of active N-glycan biosynthesis inhibition from kifunensine and swainsonine incubation in Symbiodiniaceae cells.

Swainsonine blocks MAN2, the enzyme responsible for converting hybrid glycans to complex glycans featuring even less mannose (Figure 4a). To confirm that swainsonine treatment decreased the proportion of complex glycans (and therefore increased the proportion of hybrid glycans), symbiont cells were labeled with PHA-L-PE, which binds Galβ(1-4)GlcNAc linkages from the 2,6 branch found mainly in complex glycans [71]. Symbiont cells exhibited low levels of labeling with PHA-L-PE overall, though increases in median fluorescence intensity were significant compared to unlabeled negative control (Figure 4e; ANOVA, *p <* 1 x 10^-7^). Opposite to the expectation that swainsonine blocks complex glycan formation, cells treated with swainsonine showed increased, rather than decreased PHA-L-PE labeling compared to untreated cells (Figure 4e; ANOVA, *p <* 1 x 10^-7^). These significant differences were also present in geometric and arithmetic mean fluorescence intensity comparisons (ANOVA, *p <* 1 x 10^-7^). The low levels of staining combined with an absence of complex glycan inhibition suggests differences in later steps of N-glycan biosynthesis among Symbiodiniaceae when compared to the canonical pathway. It is possible that *Breviolum minutum* has alternative pathways to build N-glycans besides the late stage processing in the Golgi apparatus, known in eukaryotes to create N-glycoproteins that can be recognized by PHA-L. Additionally, since the MAN2 found in *B. minutum* is most similar to lysosomal mannosidases (Supplementary Table 1, Supplementary Figure 2), swainsonine may have inhibited lysosomal catabolism, contributing to glycan dysregulation [72]. It is also possible that while the treatment failed to decrease complex glycan proportion, it nevertheless may have increased hybrid glycan proportion, as we did not measure hybrid glycans directly. This outcome would be consistent with the elevated mannose-binding we observed directly with CVN-PE in swainsonine-treated cells.

Kifunensine-treated cells were also expected to have a lower proportion of complex glycans, however kifunensine showed no difference in PHA-L-PE labeling compared to untreated cells (Figure 4e; ANOVA, *p* = 0.29). This lack of significant change in complex N-glycan abundance may indicate that the naturally occurring glycans on the Symbiodiniaceae cell surface are not primarily composed of complex N-glycans. Earlier studies on Symbiodiniaceae cell surface glycans found strong PHA-L lectin binding in the family [64]. However, compared to other genera, *B. minutum* exhibited the least amount of binding from PHA-L and ConA, recognizing complex N-glycans and high-mannose glycans, respectively. It is therefore possible that *B. minutum* does not have the ability to produce complex N-glycans through its own N-glycan biosynthesis. However, the presence of additional N-glycan transferases for moieties such as xylose, fucose, and galactose, as well as the detection of these and other ornaments via LC-MS/MS, support the potential for glycan complexity. It cannot be ruled out that *B. minutum* and other Symbiodiniaceae have alternate pathways for producing complex glycans. The O-glycan pathway is currently uncharacterized in Symbiodiniaceae, but it may allow for similar variation in complexity and subsequently play a role in host recognition.

### N-Glycan Biosynthesis Inhibition Decreases Symbiont Colonization Success in Host Anemones

Finally, we inoculated aposymbiotic Aiptasia polyps with *Breviolum minutum* cultures treated with the N-glycan biosynthesis inhibitors to determine whether glycan composition has an effect on symbiont colonization rates in naïve hosts. After a fixed exposure period, Aiptasia polyps inoculated with kifunensine-treated cells had a lower symbiont density compared to those inoculated with untreated control cells (Figure 5; ANOVA, Dunnett’s test, *p <* 0.05). However, animals colonized by swainsonine-treated cells had symbiont densities equivalent to controls (Figure 5; ANOVA, Dunnett’s test *p* = 0.295).

**Figure 5.**
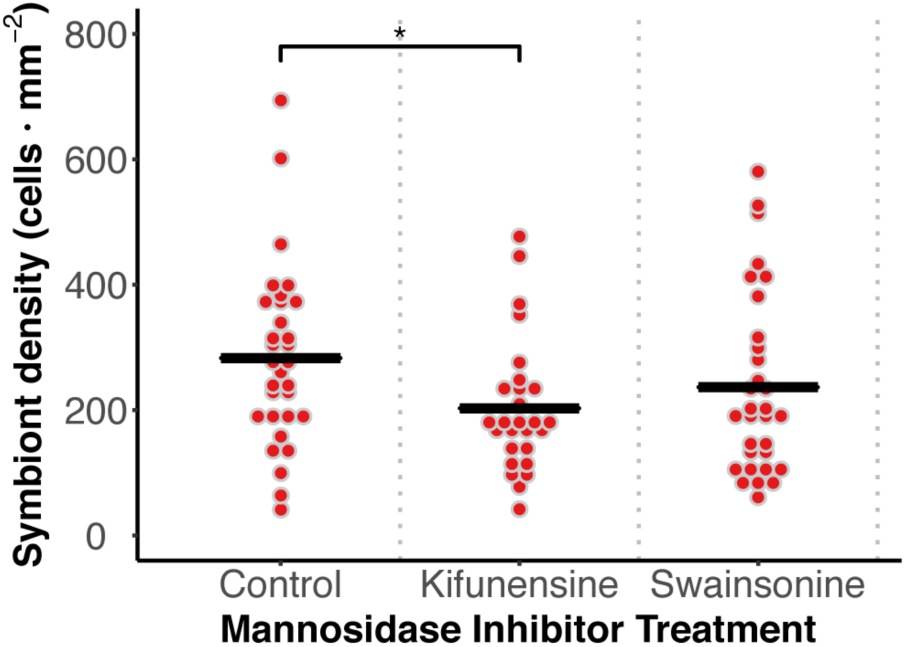
Effect of kifunensine and swainsonine symbiont colonization success of host anemones. Effect of N-glycan biosynthesis inhibition on colonization by *B. minutum* in aposymbiotic adult Aiptasia polyps (n = 30). Asterisks indicate a significant difference between treatments (ANOVA, Dunnett’s test, *p* < 0.05).

While lectin masking and enzymatic cleavage experiments have shown that low abundances of high-mannose glycans on symbiont cell surfaces reduce colonization [26–29], our biochemical inhibition data are the first to indicate that elevated proportions of high-mannose glycans can also reduce colonization, suggesting that glycan composition is a key factor modulating symbiosis establishment. Unless the proportion of different glycans is within a specific range, the micro-alga may not be recognized as an appropriate symbiont, providing a potential mechanism for maintaining specificity. An additional possibility is that interactions between lectins and hybrid glycans are selective at a later step after initial recognition of mannose. This would be consistent with the observed difference in colonization success between swainsonine-treated cells (i.e. more high-mannose and hybrid N-glycans) and kifunensine-treated cells (i.e. more high-mannose N-glycans only). A multi-step glycan recognition pathway could be part of a “winnowing” process of host-symbiont selection similar to that found in other symbioses [73].

In cnidarians, mannose recognition includes a variety of highly conserved mechanisms [74–76]. For example, the lectin-complement pathway is functional in the gastroderm and expression is upregulated in response to bacterial pathogens and algal symbionts [74, 77]. The binding of pattern recognition receptors, such as the coral mannose-binding lectin millectin, to both pathogenic bacteria and members of Symbiodiniaceae suggests a dual role of mannose glycan recognition in cnidarian immunity and symbiosis [17, 78]. Indeed, the activation of host innate immunity during colonization contrasts with the repression of host innate immunity under stable symbiosis conditions [77, 79–81]. Therefore, the response of host cells to symbiont N-glycan abundance and composition may differ depending on symbiotic state and life history stage.

### Implications and Future Directions

Our results provide new information about the chemical signaling molecules of *Breviolum minutum*, a mutualistic dinoflagellate symbiont of corals and sea anemones. We verified that the N-linked glycan biosynthesis pathway is mostly present and conserved in the *B. minutum* genome, and we determined the structure and abundance of N-glycans cleaved from the algal surface. While our experiments stress the notion that N-glycans are important for symbiont recognition, we observed a complex algal glycome without clear determinants of symbiosis establishment. The direct approach of manipulating this algal glycome (through cleavage with PNGase F or through biosynthesis inhibition) has had a more consistent effect on colonization than glycan masking approaches [26, 28, 29, 57]. Nevertheless, colonization success was highly variable in the clonal adult hosts that were used in this study, possibly due to small differences in polyp size or states of innate immunity in aposymbiotic adults. Challenging naïve, aposymbiotic larvae may be a more fruitful approach in the Aiptasia system when investigating symbiosis establishment, since larvae allow for greater sample sizes and thus more statistical power to detect subtle differences in colonization [57]. Despite this limitation, the differences observed in glycan characterization and symbiont recolonization support the further use of aposymbiotic Aiptasia adults as a model for understanding processes involved in bleaching recovery. The potential for variation in N-glycan biosynthesis among Symbiodiniaceae could provide a basis for host-symbiont recognition as well as recolonization dynamics after environmental conditions that result in dysbiosis.

## 2.5 Acknowledgements

This research was funded by the National Science Foundation (NSF IOS-1557804 to V.M.W and S.L.). We thank Shumpei Maruyama, Sarah Howey, Milan Sengthrep, Tyler Coleman, Kyle Petersen, Darian Thompson, and Kenneth Moller for their assistance in algal and animal maintenance. In addition, we thank Allison Ehrlich, Jamie Pennington, and the Environmental and Molecular Toxicology Department at Oregon State University for technical support in flow cytometry.

## SUPPLEMENTARY MATERIAL

**Supplementary Figure 1.**
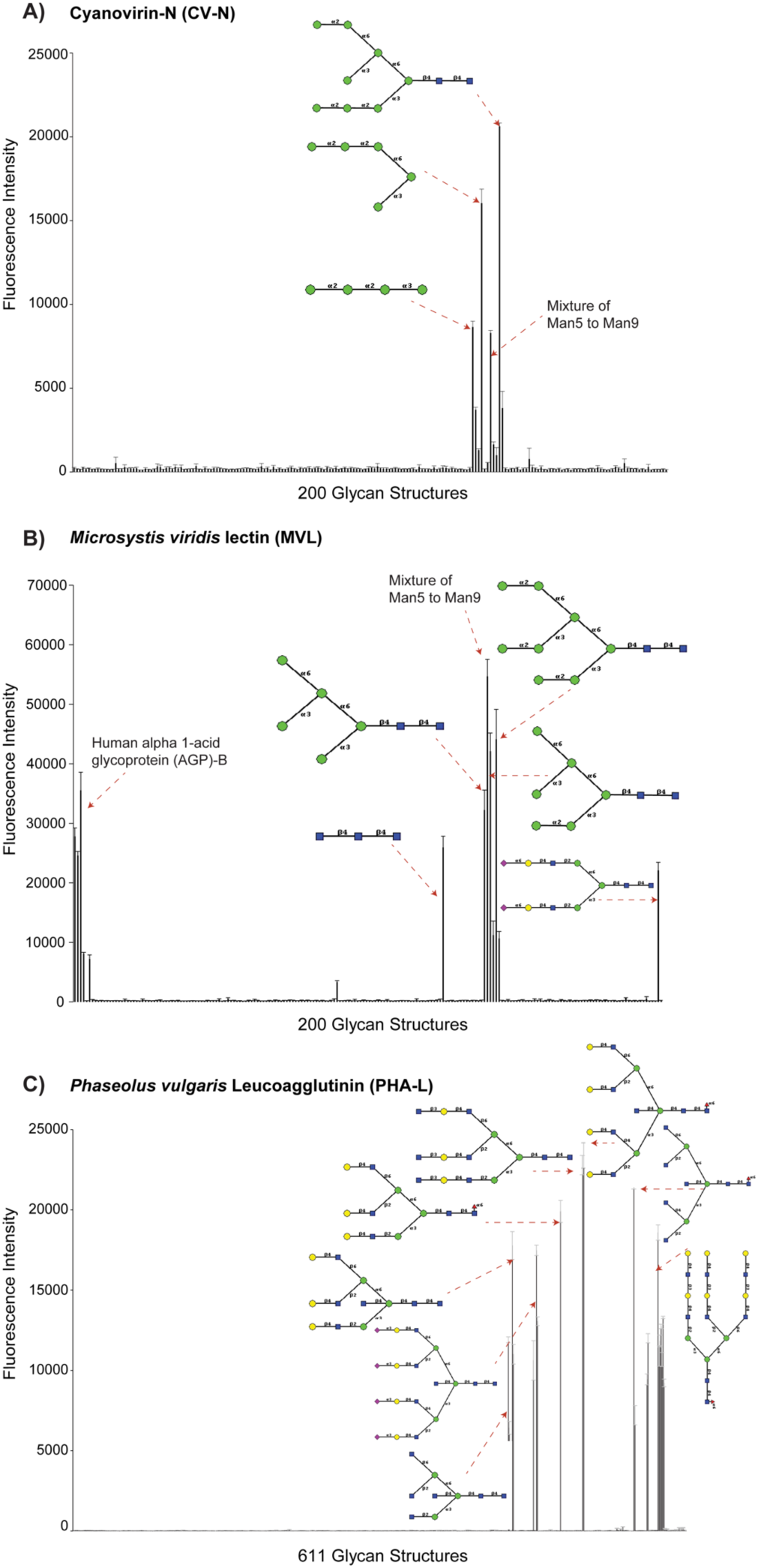
Lectin specificity of MVL, CVN, and PHA-L. The lectin specificity of (**a**) MVL, (**b**) CVN, and (**c**) PHA-L. is shown as determined from glycan microarray experiments. MVL shows binding affinity for chitobiose core N-glycan motifs [Man-3 to Man-9; α(1-6) and α(1-3)-dimannoside, Man β(1-4) GlcNAc], CVN shows affinity for high-mannose N-glycan motifs [Man-8 and Man-9 only; α(1-2)-dimannoside], and PHA-L shows affinity for complex type N-glycan motifs [Gal-β(1-4)GlcNAc β(1-6)(GlcNAc β(1-2)Man α(1-3)) Manα(1-3)]. Data provided by the Consortium for Functional Glycomics.

**Supplementary Figure 2.**
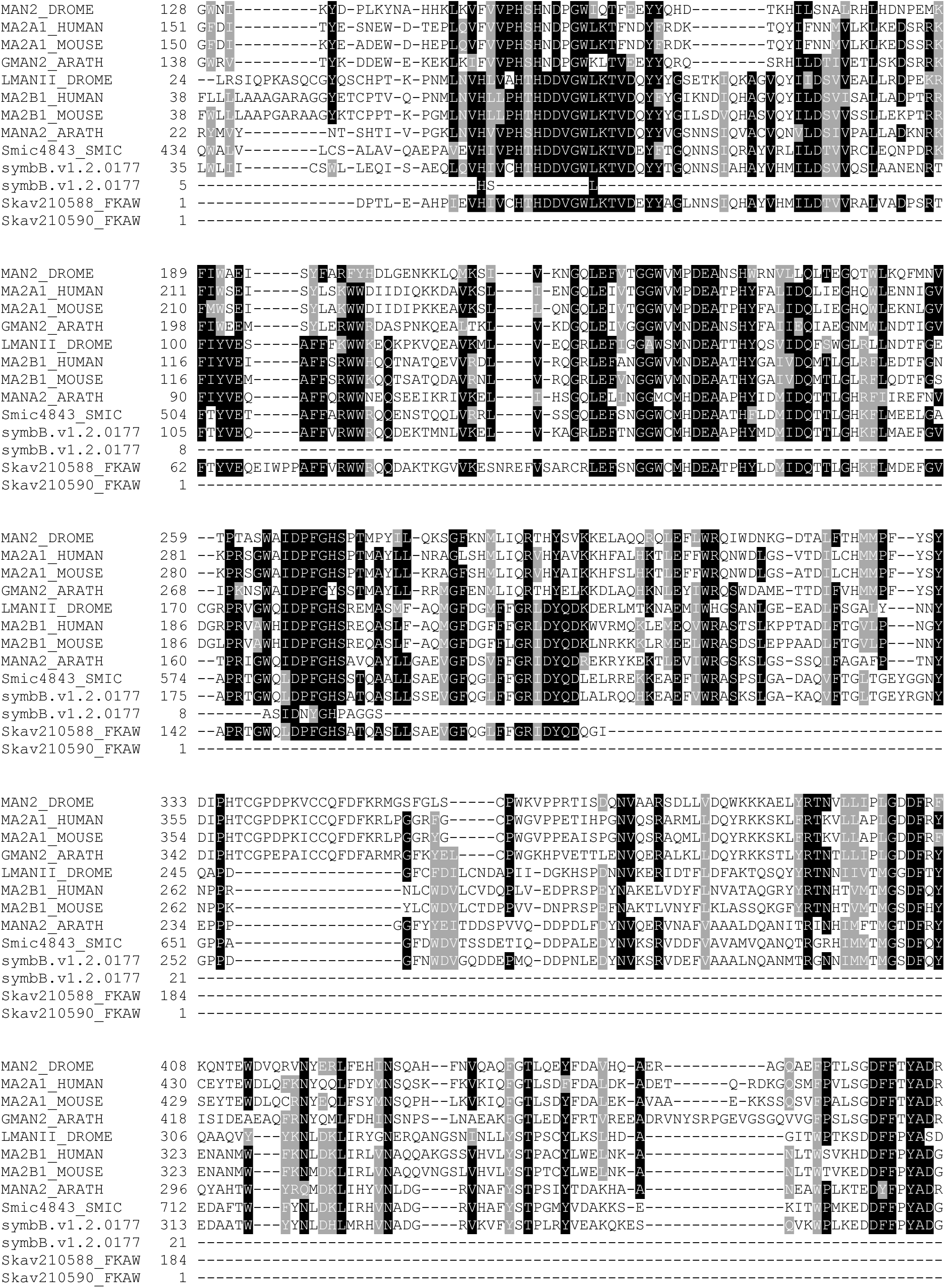
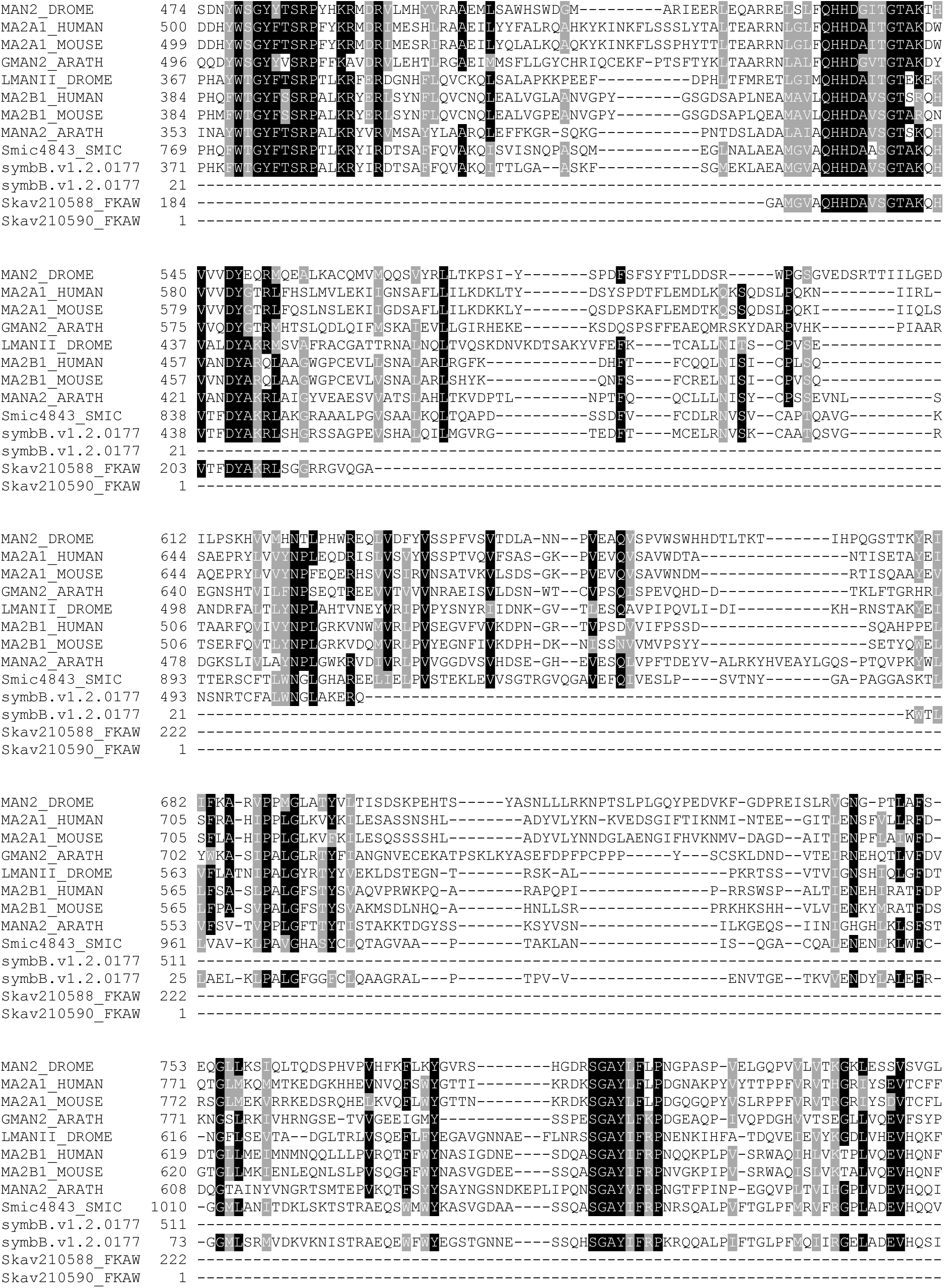
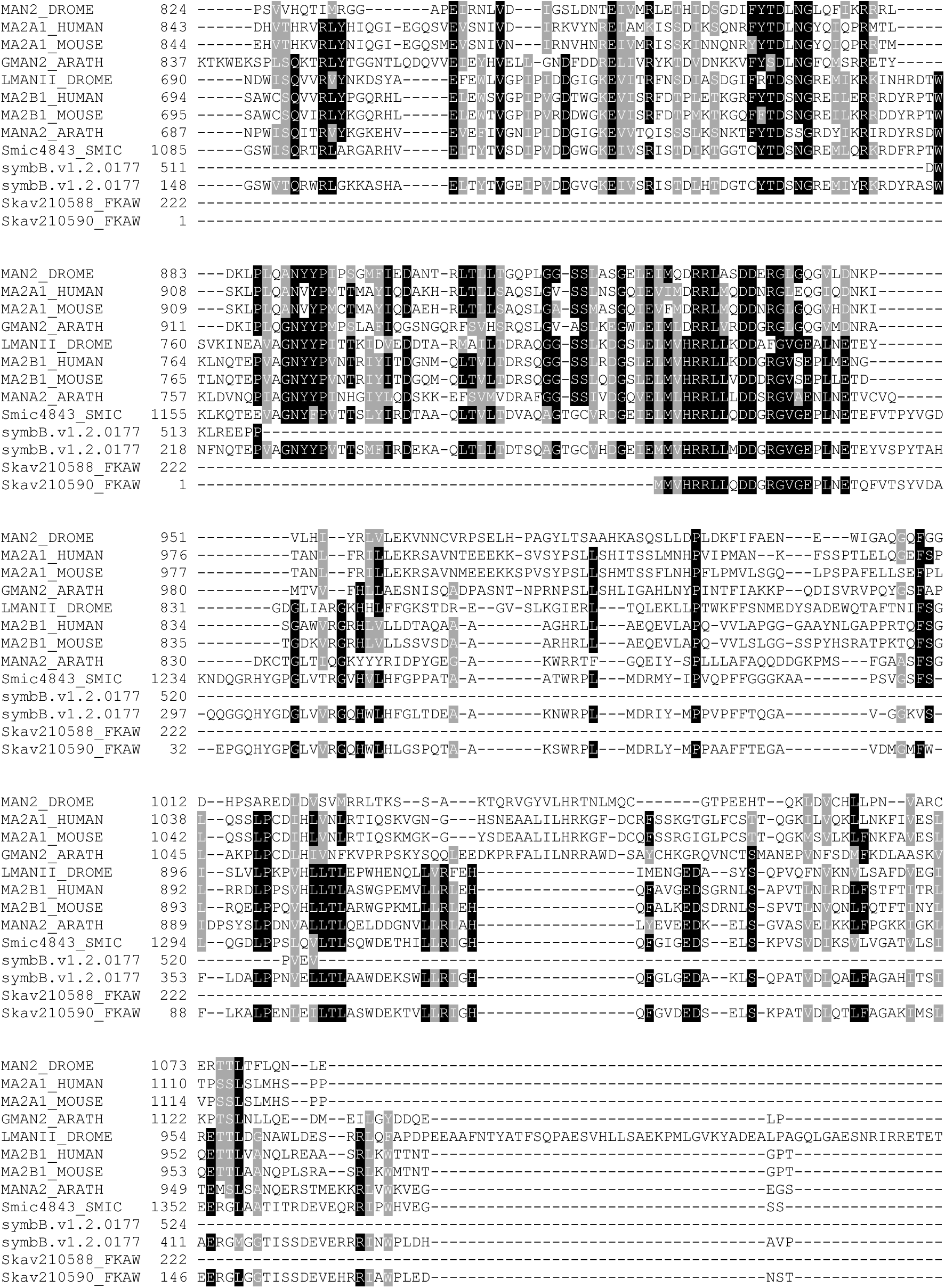
Multiple sequence alignment of Mannosidase II sequences. A multiple sequence alignment was conducted in T-Coffee using the amino acid sequences of Golgi Mannosidase II (MAN2_DROME, MA2A1_HUMAN, MA2A1_MOUSE, GMAN2_ARATH) and Lysosomal α-Mannosidase (LMANII_DROME, MA2B1_HUMAN, MA2B1_MOUSE, MANA2_ARATH) from *Drosophila melanogaster*, *Homo sapiens*, *Mus musculus*, and *Arabidopsis thaliana* to the α-Mannosidase found in Symbiodiniaceae species *Symbiodinium microadriaticum* (Smic), *Breviolum minutum (*symbB), and *Fugacium kawagutii* (Fkaw).

**Supplementary Table 1:**
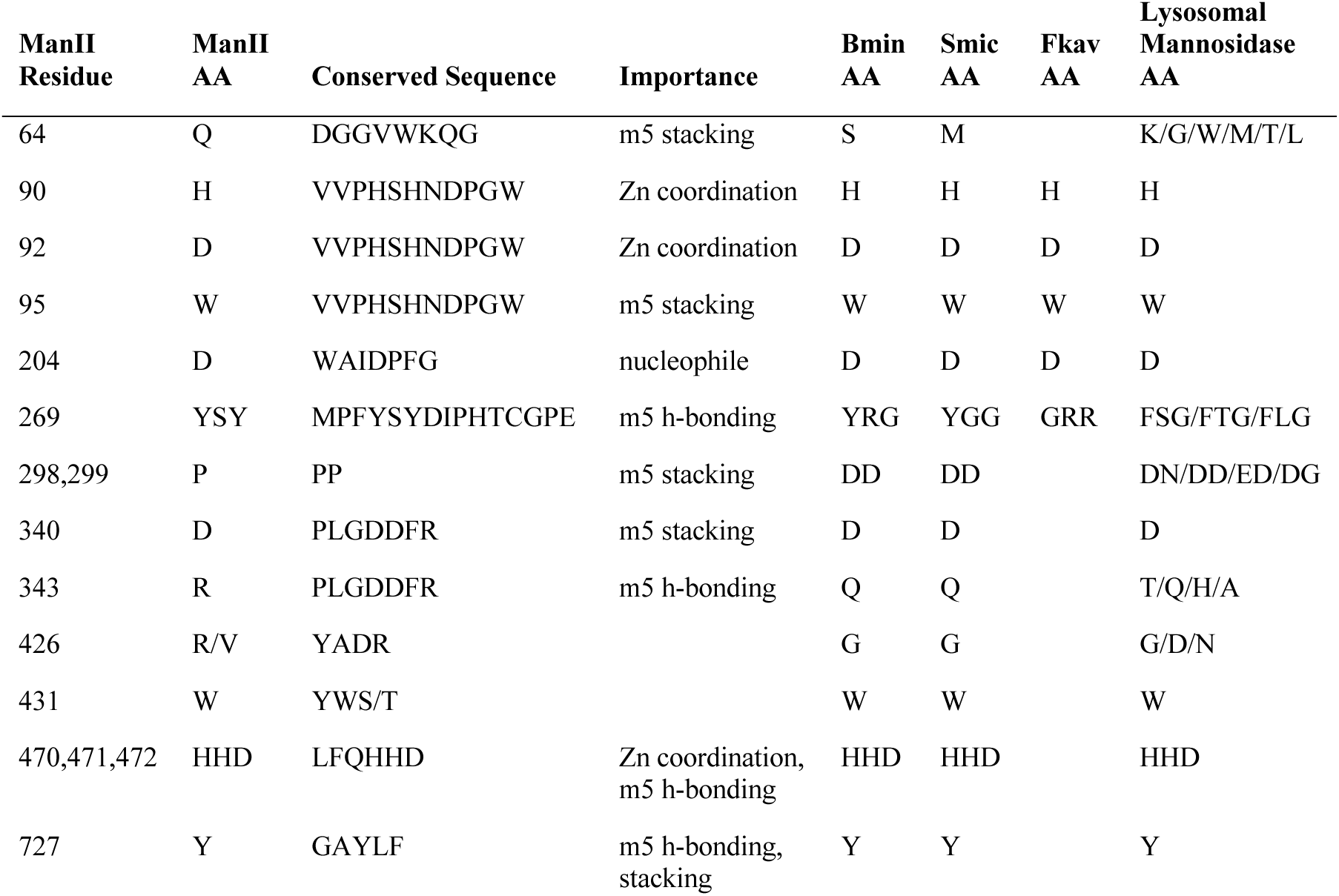
ManII site conservation in Symbiodiniaceae. The conserved ManII residue in *Drosophila* is shown on the left, with its corresponding amino acid and conserved sequence. The importance of the residue for binding affinity is shown in the center. Symbiodiniaceae residues are shown for *B. minutum*, *S. microadriaticum*, and *F. kawagutii.* Conserved residues for lysosomal mannosidase amino acids are shown on the right.

**Supplementary Table 2:**
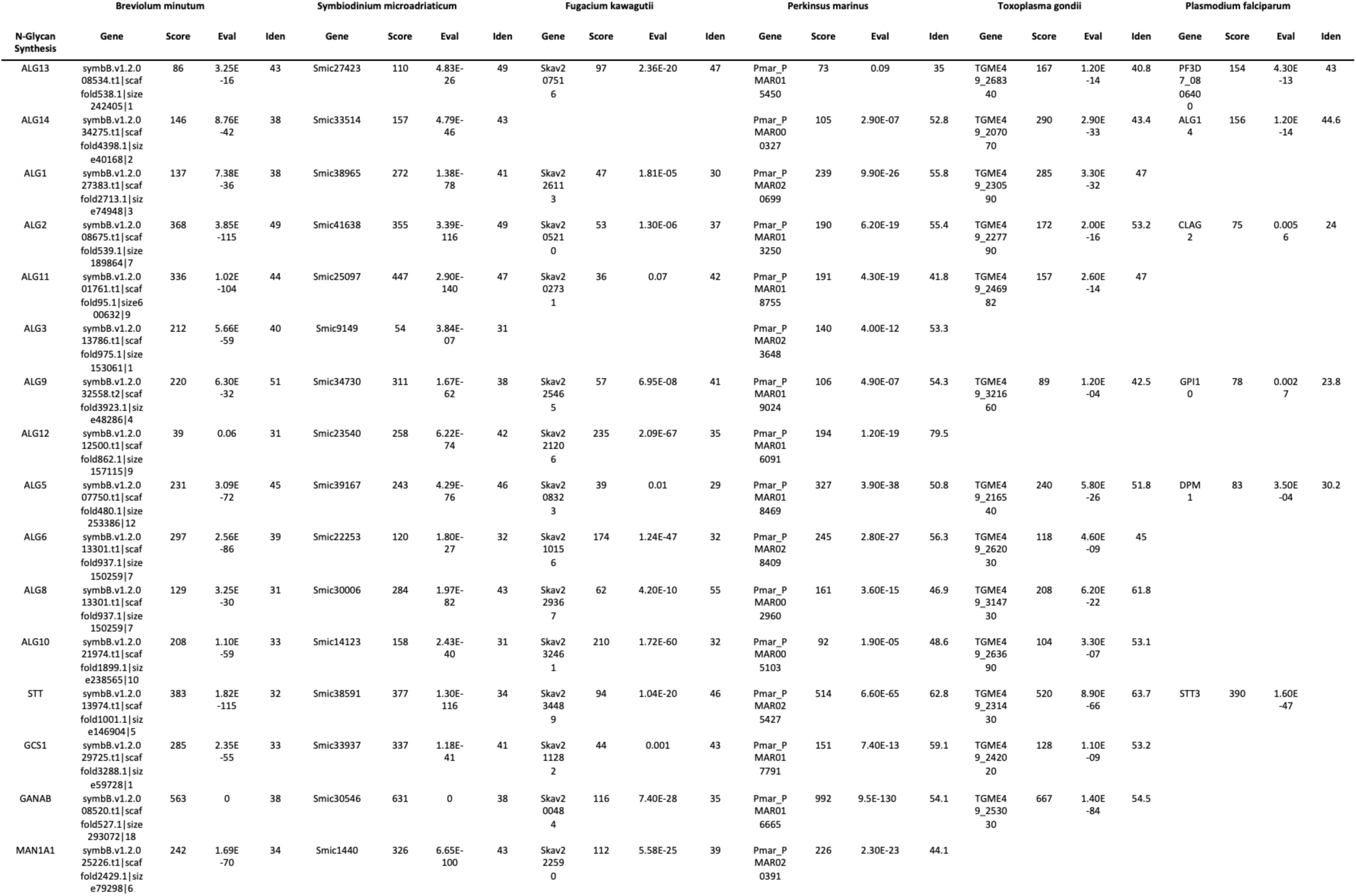
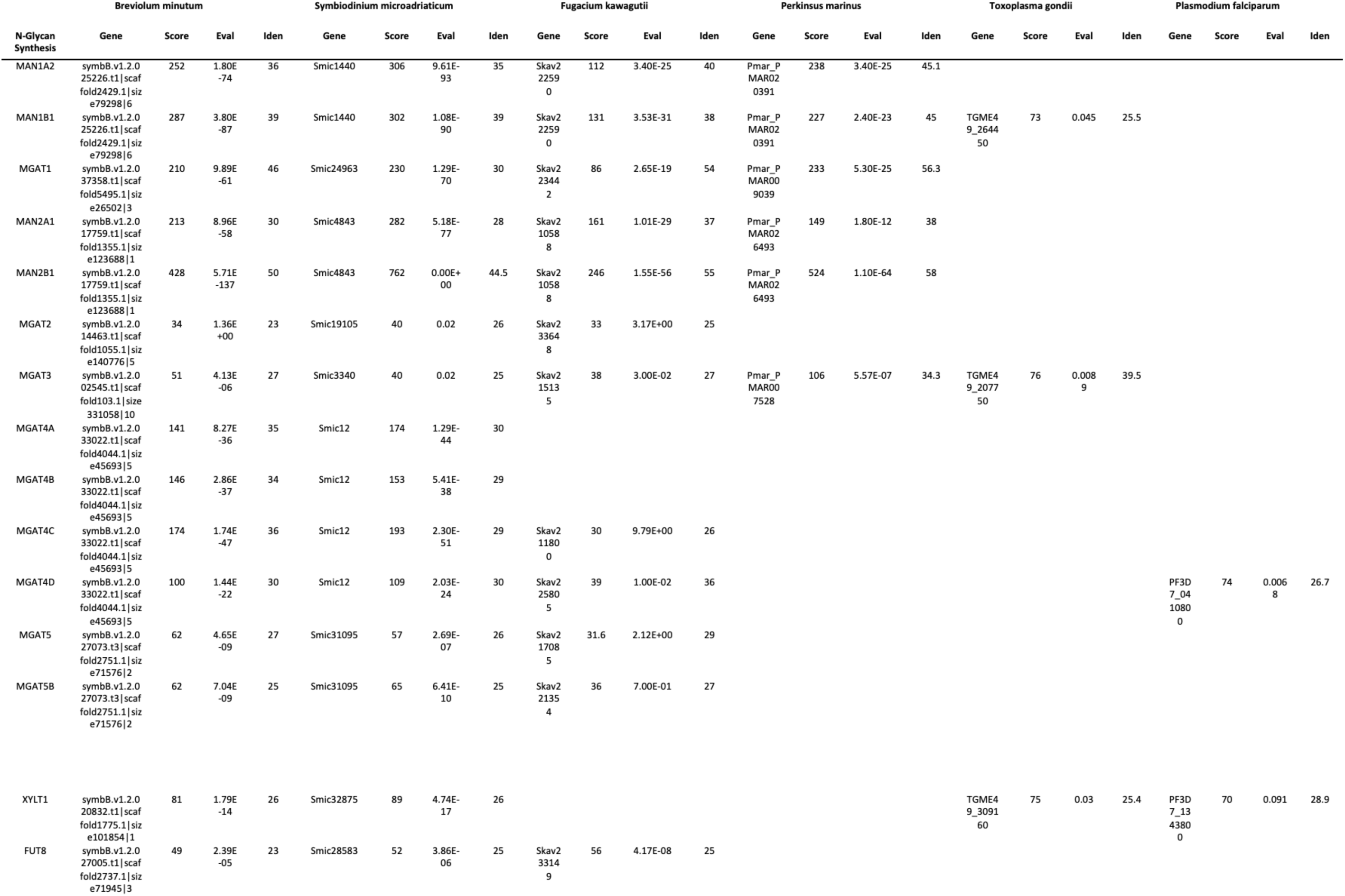
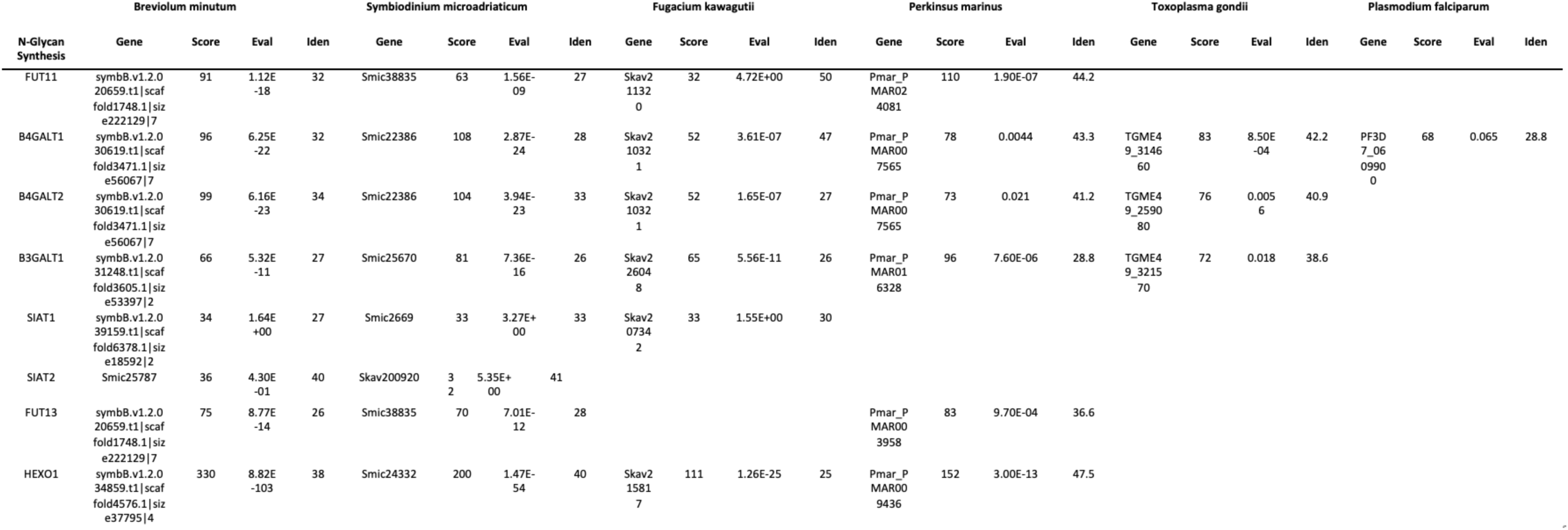
N-glycan biosynthesis datasheet. N-glycan biosynthesis BLASTp searches were performed in Symbiodiniaceae species and in related alveolates *Perkinsus marinus*, *Toxoplasma gondii*, and *Plasmodium falciparum*. Manually annotated N-glycan protein sequences from humans were collected from Uniprot and queried directly against a target species protein database consisting of gene models. The first result of each successful search (gene) is shown for each protein query, along with its score, eval, and % identity.

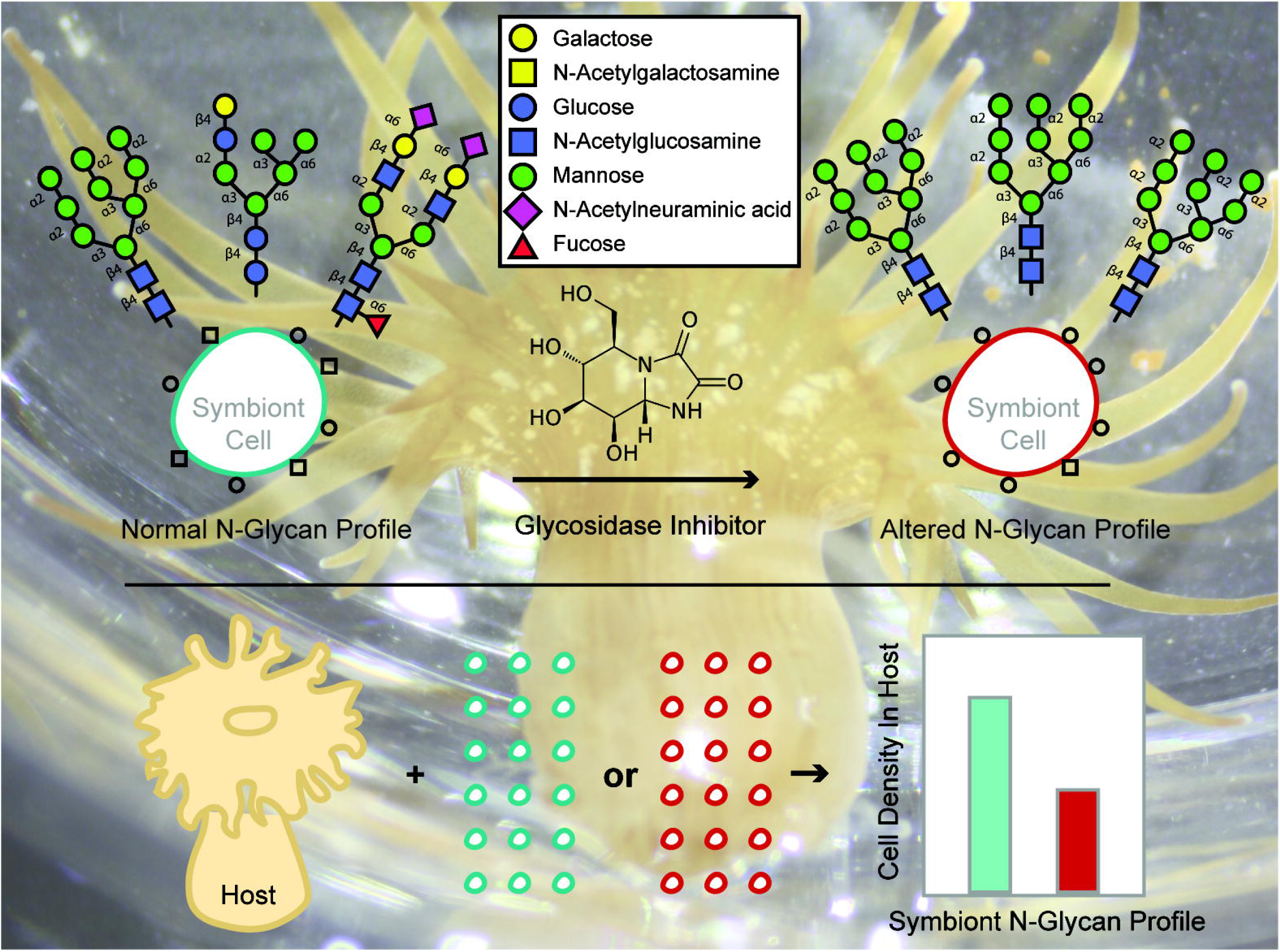

## References

1. LaJeunesse TC, Parkinson JE, Gabrielson PW, et al (2018) Systematic Revision of Symbiodiniaceae Highlights the Antiquity and Diversity of Coral Endosymbionts. Current Biology 28:2570–2580.e6. https://doi.org/10.1016/j.cub.2018.07.008

2. Davy SK, Allemand D, Weis VM (2012) Cell Biology of Cnidarian-Dinoflagellate Symbiosis. Microbiology and Molecular Biology Reviews 76:229–261. https://doi.org/10.1128/MMBR.05014-11

3. Weis VM (2019) Cell Biology of Coral Symbiosis: Foundational Study Can Inform Solutions to the Coral Reef Crisis. Integrative and Comparative Biology. https://doi.org/10.1093/icb/icz067

4. Bosch TCG (2013) Cnidarian-microbe interactions and the origin of innate immunity in metazoans. Annual review of microbiology 67:499–518. https://doi.org/10.1146/annurev-micro-092412-155626

5. Dunn SR (2009) Immunorecognition and immunoreceptors in the Cnidaria. Invertebrate Survival Journal 6:7–14

6. Hamada M, Shoguchi E, Shinzato C, et al (2013) The complex NOD-like receptor repertoire of the coral Acropora digitifera includes novel domain combinations. Molecular biology and evolution 30:167–76. https://doi.org/10.1093/molbev/mss213

7. Neubauer E-F, Poole AZ, Neubauer P, et al (2017) A diverse host thrombospondin-type-1 repeat protein repertoire promotes symbiont colonization during establishment of cnidarian-dinoflagellate symbiosis. eLife 6:. https://doi.org/10.7554/eLife.24494

8. Poole AZ, Weis VM (2014) TIR-domain-containing protein repertoire of nine anthozoan species reveals coral-specific expansions and uncharacterized proteins. Developmental and Comparative Immunology 46:480–488. https://doi.org/10.1016/j.dci.2014.06.002

9. McGuinness DH, Dehal PK, Pleass RJ (2003) Pattern recognition molecules and innate immunity to parasites. Trends in Parasitology 19:312–319. https://doi.org/10.1016/S1471-4922(03)00123-5

10. Varki A (2017) Biological roles of glycans. Glycobiology 27:3–49. https://doi.org/10.1093/glycob/cww086

11. Collins BE, Paulson JC (2004) Cell surface biology mediated by low affinity multivalent protein–glycan interactions. Current Opinion in Chemical Biology 8:617–625. https://doi.org/10.1016/j.cbpa.2004.10.004

12. Gallatin WM, Weissman IL, Butcher EC (1983) A cell-surface molecule involved in organ-specific homing of lymphocytes. Nature 304:30–34. https://doi.org/10.1038/304030a0

13. Pang P-C, Chiu PCN, Lee C-L, et al (2011) Human Sperm Binding Is Mediated by the Sialyl-Lewisx Oligosaccharide on the Zona Pellucida. Science 333:1761–1764. https://doi.org/10.1126/science.1207438

14. Lannoo N, Van Damme EJM (2014) Lectin domains at the frontiers of plant defense. Front Plant Sci 5:. https://doi.org/10.3389/fpls.2014.00397

15. Baum LG, Garner OB, Schaefer K, Lee B (2014) Microbe-host interactions are positively and negatively regulated by galectin-glycan interactions. Frontiers in Immunology 5:1–8. https://doi.org/10.3389/fimmu.2014.00284

16. Kachko A, Loesgen S, Shahzad-Ul-Hussan S, et al (2013) Inhibition of hepatitis C virus by the cyanobacterial protein Microcystis viridis lectin: mechanistic differences between the high-mannose specific lectins MVL, CV-N, and GNA. Molecular pharmaceutics 10:4590– 602. https://doi.org/10.1021/mp400399b

17. Kvennefors ECE, Leggat W, Hoegh-Guldberg O, et al (2008) An ancient and variable mannose-binding lectin from the coral Acropora millepora binds both pathogens and symbionts. Developmental & Comparative Immunology 32:1582–1592. https://doi.org/10.1016/j.dci.2008.05.010

18. Zhou Z, Yu X, Tang J, et al (2017) Dual recognition activity of a rhamnose-binding lectin to pathogenic bacteria and zooxanthellae in stony coral Pocillopora damicornis. Developmental & Comparative Immunology 70:88–93. https://doi.org/10.1016/j.dci.2017.01.009

19. Ip WK, Takahashi K, Alan Ezekowitz R, Stuart LM (2009) Mannose-binding lectin and innate immunity. Immunological Reviews 230:9–21. https://doi.org/10.1111/j.1600-065X.2009.00789.x

20. van Kooyk Y, Rabinovich G a (2008) Protein-glycan interactions in the control of innate and adaptive immune responses. Nature immunology 9:593–601. https://doi.org/10.1038/ni.f.203

21. Schwarz JA (2008) Understanding the intracellular niche in cnidarian-Symbiodinium symbioses: parasites lead the way. Vie et Milieu - Life and Environment 58:141–151

22. Bushkin GG, Ratner DM, Cui J, et al (2010) Suggestive evidence for Darwinian selection against asparagine-linked glycans of Plasmodium falciparum and Toxoplasma gondii. Eukaryotic Cell 9:228–241. https://doi.org/10.1128/EC.00197-09

23. Fauquenoy S, Morelle W, Hovasse A, et al (2008) Proteomics and Glycomics Analyses of N-Glycosylated Structures Involved in Toxoplasma gondii-Host Cell Interactions. Molecular & Cellular Proteomics 7:891–910. https://doi.org/10.1074/mcp.M700391-MCP200

24. Samuelson J, Banerjee S, Magnelli P, et al (2005) The diversity of dolichol-linked precursors to Asn-linked glycans likely results from secondary loss of sets of glycosyltransferases. Proceedings of the National Academy of Sciences 102:1548–1553. https://doi.org/10.1073/pnas.0409460102

25. Tasumi S, Vasta GR (2007) A Galectin of Unique Domain Organization from Hemocytes of the Eastern Oyster (Crassostrea virginica) Is a Receptor for the Protistan Parasite Perkinsus marinus. The Journal of Immunology 179:3086–3098. https://doi.org/10.4049/jimmunol.179.5.3086

26. Bay LK, Cumbo VR, Abrego D, et al (2011) Infection Dynamics Vary between Symbiodinium Types and Cell Surface Treatments during Establishment of Endosymbiosis with Coral Larvae. Diversity 3:356–374. https://doi.org/10.3390/d3030356

27. Kuniya N, Jimbo M, Tanimoto F, et al (2015) Possible involvement of Tachylectin-2-like lectin from Acropora tenuis in the process of Symbiodinium acquisition. Fisheries Science 81:473–483. https://doi.org/10.1007/s12562-015-0862-y

28. Lin K, Wang J, Fang L-S (2000) Participation of Glycoproteins on Zooxanthellal Cell Walls in the Establishment of a Symbiotic Relationship with the Sea Anemone, Aiptasia pulchella. 39:172–178

29. Wood-Charlson EM, Hollingsworth LL, Krupp DA, Weis VM (2006) Lectin/glycan interactions play a role in recognition in a coral/dinoflagellate symbiosis. Cellular Microbiology 8:1985–1993. https://doi.org/10.1111/j.1462-5822.2006.00765.x

30. Helenius A, Aebi M (2004) Roles of N-Linked Glycans in the Endoplasmic Reticulum. Annual Review of Biochemistry 73:1019–1049. https://doi.org/10.1146/annurev.biochem.73.011303.073752

31. Stanley P, Taniguchi N, Aebi M (2015) N-Glycans. In: Varki A, Cummings RD, Esko JD, et al (eds) Essentials of Glycobiology, 3rd ed. Cold Spring Harbor Laboratory Press, Cold Spring Harbor (NY)

32. Elbein AD, Tropea JE, Mitchell M, Kaushal GP (1990) Kifunensine, a potent inhibitor of the glycoprotein processing mannosidase I. J Biol Chem 265:15599–15605

33. Liebminger E, Hüttner S, Vavra U, et al (2009) Class I alpha-mannosidases are required for N-glycan processing and root development in Arabidopsis thaliana. The Plant cell 21:3850– 3867. https://doi.org/10.1105/tpc.109.072363

34. Weng S, Spiro RG (1993) Demonstration that a kifunensine-resistant alpha-mannosidase with a unique processing action on N-linked oligosaccharides occurs in rat liver endoplasmic reticulum and various cultured cells. J Biol Chem 268:25656–25663

35. Tulsiani DR, Harris TM, Touster O (1982) Swainsonine inhibits the biosynthesis of complex glycoproteins by inhibition of Golgi mannosidase II. J Biol Chem 257:7936–7939

36. Petersen L, Ardèvol A, Rovira C, Reilly PJ (2010) Molecular Mechanism of the Glycosylation Step Catalyzed by Golgi α-Mannosidase II: A QM/MM Metadynamics Investigation. Journal of the American Chemical Society 132:8291–8300. https://doi.org/10.1021/ja909249u

37. Shah N, Kuntz DA, Rose DR (2008) Golgi-mannosidase II cleaves two sugars sequentially in the same catalytic site. Proceedings of the National Academy of Sciences 105:9570– 9575. https://doi.org/10.1073/pnas.0802206105

38. Shoguchi E, Shinzato C, Kawashima T, et al (2013) Draft assembly of the Symbiodinium minutum nuclear genome reveals dinoflagellate gene structure. Current biology : CB 23:1399–408. https://doi.org/10.1016/j.cub.2013.05.062

39. Aranda M, Li Y, Liew YJ, et al (2016) Genomes of coral dinoflagellate symbionts highlight evolutionary adaptations conducive to a symbiotic lifestyle. Scientific Reports 6:39734. https://doi.org/10.1038/srep39734

40. Lin S, Cheng S, Song B, et al (2015) The Symbiodinium kawagutii genome illuminates dinoflagellate gene expression and coral symbiosis. Science 350:691–694. https://doi.org/10.1126/science.aad0408

41. Joseph SJ, Fernández-Robledo JA, Gardner MJ, et al (2010) The Alveolate Perkinsus marinus: Biological Insights from EST Gene Discovery. BMC Genomics 11:228. https://doi.org/10.1186/1471-2164-11-228

42. Gardner MJ, Hall N, Fung E, et al (2002) Genome sequence of the human malaria parasite Plasmodium falciparum. Nature 419:. https://doi.org/10.1038/nature01097

43. Kissinger JC, Gajria B, Li L, et al (2003) ToxoDB: accessing the Toxoplasma gondii genome. Nucleic Acids Res 31:234–236. https://doi.org/10.1093/nar/gkg072

44. The UniProt Consortium (2017) UniProt: the universal protein knowledgebase. Nucleic Acids Research 45:D158–D169. https://doi.org/10.1093/nar/gkw1099

45. Liew YJ, Aranda M, Voolstra CR (2016) Reefgenomics.Org - a repository for marine genomics data. Database (Oxford) 2016:. https://doi.org/10.1093/database/baw152

46. Kersey PJ, Allen JE, Allot A, et al (2018) Ensembl Genomes 2018: an integrated omics infrastructure for non-vertebrate species. Nucleic Acids Res 46:D802–D808. https://doi.org/10.1093/nar/gkx1011

47. Di Tommaso P, Moretti S, Xenarios I, et al (2011) T-Coffee: a web server for the multiple sequence alignment of protein and RNA sequences using structural information and homology extension. Nucleic Acids Res 39:W13–W17. https://doi.org/10.1093/nar/gkr245

48. Notredame C, Higgins DG, Heringa J (2000) T-Coffee: A novel method for fast and accurate multiple sequence alignment. J Mol Biol 302:205–217. https://doi.org/10.1006/jmbi.2000.4042

49. Park C, Meng L, Stanton LH, et al (2005) Characterization of a Human Core-specific Lysosomal α1,6-Mannosidase Involved in *N*-Glycan Catabolism. Journal of Biological Chemistry 280:37204–37216. https://doi.org/10.1074/jbc.M508930200

50. Dong X, Zhou S, Mechref Y (2016) LC-MS/MS analysis of permethylated free oligosaccharides and N-glycans derived from human, bovine, and goat milk samples. ELECTROPHORESIS 37:1532–1548. https://doi.org/10.1002/elps.201500561

51. Kang P, Mechref Y, Novotny MV (2008) High-throughput solid-phase permethylation of glycans prior to mass spectrometry. Rapid Communications in Mass Spectrometry 22:721– 734. https://doi.org/10.1002/rcm.3395

52. Zhong J, Banazadeh A, Peng W, Mechref Y (2018) A carbon nanoparticles-based solid-phase purification method facilitating sensitive MALDI–MS analysis of permethylated N-glycans. ELECTROPHORESIS 39:3087–3095. https://doi.org/10.1002/elps.201800254

53. Zhu R, Zhou S, Peng W, et al (2018) Enhanced Quantitative LC-MS/MS Analysis of N-linked Glycans Derived from Glycoproteins Using Sodium Deoxycholate Detergent. J Proteome Res 17:2668–2678. https://doi.org/10.1021/acs.jproteome.8b00127

54. Yu C-Y, Mayampurath A, Hu Y, et al (2013) Automated annotation and quantification of glycans using liquid chromatography–mass spectrometry. Bioinformatics 29:1706–1707. https://doi.org/10.1093/bioinformatics/btt190

55. Shahzad-ul-Hussan S, Cai M, Bewley CA (2009) Unprecedented Glycosidase Activity at a Lectin Carbohydrate-Binding Site Exemplified by the Cyanobacterial Lectin MVL. https://pubs.acs.org/doi/full/10.1021/ja905929c. Accessed 13 Feb 2019

56. Matthews JL, Sproles AE, Oakley CA, et al (2016) Menthol-induced bleaching rapidly and effectively provides experimental aposymbiotic sea anemones (Aiptasia sp.) for symbiosis investigations. Journal of Experimental Biology 219:306–310. https://doi.org/10.1242/jeb.128934

57. Parkinson JE, Tivey TR, Mandelare PE, et al (2018) Subtle Differences in Symbiont Cell Surface Glycan Profiles Do Not Explain Species-Specific Colonization Rates in a Model Cnidarian-Algal Symbiosis. Frontiers in Microbiology 9:. https://doi.org/10.3389/fmicb.2018.00842

58. Baumgarten S, Simakov O, Esherick LY, et al (2015) The genome of Aiptasia, a sea anemone model for coral symbiosis. Proceedings of the National Academy of Sciences 112:11893–11898. https://doi.org/10.1073/pnas.1513318112

59. Sunagawa S, Wilson EC, Thaler M, et al (2009) Generation and analysis of transcriptomic resources for a model system on the rise: the sea anemone Aiptasia pallida and its dinoflagellate endosymbiont. BMC Genomics 10:258. https://doi.org/10.1186/1471-2164-10-258

60. Liu H, Stephens TG, González-Pech RA, et al (2018) Symbiodinium genomes reveal adaptive evolution of functions related to coral-dinoflagellate symbiosis. Communications Biology 1:. https://doi.org/10.1038/s42003-018-0098-3

61. Davidson EA, Gowda DC (2001) Glycobiology of Plasmodium falciparum. Biochimie 83:601–604. https://doi.org/10.1016/S0300-9084(01)01316-5

62. von Itzstein M, Plebanski M, Cooke BM, Coppel RL (2008) Hot, sweet and sticky: the glycobiology of Plasmodium falciparum. Trends in Parasitology 24:210–218. https://doi.org/10.1016/j.pt.2008.02.007

63. Zhou S, Dong X, Veillon L, et al (2017) LC-MS/MS analysis of permethylated N-glycans facilitating isomeric characterization. Analytical and Bioanalytical Chemistry 409:453–466. https://doi.org/10.1007/s00216-016-9996-8

64. Logan DDK, LaFlamme AC, Weis VM, Davy SK (2010) Flow-Cytometric Characterization of the Cell-Surface Glycans of Symbiotic Dinoflagellates (Symbiodinium Spp.). Journal of Phycology 46:525–533. https://doi.org/10.1111/j.1529-8817.2010.00819.x

65. Varki A (2008) Sialic acids in human health and disease. Trends in Molecular Medicine 14:351–360. https://doi.org/10.1016/j.molmed.2008.06.002

66. Mamedov T, Yusibov V (2011) Green algae *Chlamydomonas reinhardtii* possess endogenous sialylated N-glycans. FEBS Open Bio 1:15–22. https://doi.org/10.1016/j.fob.2011.10.003

67. Stephens TG, Ragan MA, Bhattacharya D, Chan CX (2018) Core genes in diverse dinoflagellate lineages include a wealth of conserved dark genes with unknown functions. Scientific Reports 8:. https://doi.org/10.1038/s41598-018-35620-z

68. Warren L (1963) The distribution of sialic acids in nature. Comparative Biochemistry and Physiology 10:153–171. https://doi.org/10.1016/0010-406X(63)90238-X

69. Jimbo M, Suda Y, Koike K, et al (2013) Possible involvement of glycolipids in lectin-mediated cellular transformation of symbiotic microalgae in corals. Journal of Experimental Marine Biology and Ecology 439:129–135. https://doi.org/10.1016/j.jembe.2012.10.022

70. Kita A, Jimbo M, Sakai R, et al (2015) Crystal structure of a symbiosis-related lectin from octocoral. Glycobiology 25:1016–1023. https://doi.org/10.1093/glycob/cwv033

71. Fabre C, Causse H, Mourey L, et al (1998) Characterization and sugar-binding properties of arcelin-1, an insecticidal lectin-like protein isolated from kidney bean (Phaseolus vulgaris L. cv. RAZ-2) seeds. Biochem J 329:551–560

72. Dorling PR, Huxtable CR, Colegate SM (1980) Inhibition of lysosomal alpha-mannosidase by swainsonine, an indolizidine alkaloid isolated from Swainsona canescens. The Biochemical journal 191:649–651. https://doi.org/10.1042/BJ1910649

73. Nyholm SV, McFall-Ngai MJ (2004) The winnowing: establishing the squid-vibrio symbiosis. Nature reviews Microbiology 2:632–42. https://doi.org/10.1038/nrmicro957

74. Kimura A, Sakaguchi E, Nonaka M (2009) Multi-component complement system of Cnidaria: C3, Bf, and MASP genes expressed in the endodermal tissues of a sea anemone, Nematostella vectensis. Immunobiology 214:165–178. https://doi.org/10.1016/j.imbio.2009.01.003

75. Meyer E, Weis VM (2012) Study of cnidarian-algal symbiosis in the “omics” age. The Biological Bulletin 223:44–65. https://doi.org/10.1086/BBLv223n1p44

76. Wood-Charlson EM, Weis VM (2009) The diversity of C-type lectins in the genome of a basal metazoan, Nematostella vectensis. Developmental and comparative immunology 33:881–9. https://doi.org/10.1016/j.dci.2009.01.008

77. Poole AZ, Kitchen SA, Weis VM (2016) The Role of Complement in Cnidarian-Dinoflagellate Symbiosis and Immune Challenge in the Sea Anemone Aiptasia pallida. Frontiers in Microbiology 7:519. https://doi.org/10.3389/fmicb.2016.00519

78. Kvennefors ECE, Leggat W, Kerr CC, et al (2010) Analysis of evolutionarily conserved innate immune components in coral links immunity and symbiosis. Developmental and comparative immunology 34:1219–29. https://doi.org/10.1016/j.dci.2010.06.016

79. Detournay O, Schnitzler CE, Poole A, Weis VM (2012) Regulation of cnidarian– dinoflagellate mutualisms: Evidence that activation of a host TGFB innate immune pathway promotes tolerance of the symbiont. Developmental and Comparative Immunology 38:525–537. https://doi.org/10.1016/j.dci.2012.08.008

80. Mansfield KM, Carter NM, Nguyen L, et al (2017) Transcription factor NF-κB is modulated by symbiotic status in a sea anemone model of cnidarian bleaching. Scientific Reports 7:1– 14. https://doi.org/10.1038/s41598-017-16168-w

81. Merselis DG, Lirman D, Rodriguez-Lanetty M (2018) Symbiotic immuno-suppression: is disease susceptibility the price of bleaching resistance? PeerJ 6:e4494. https://doi.org/10.7717/peerj.4494

82. Helenius A, Aebi M (2001) Intracellular Functions of N-Linked Glycans. Science 291:2364– 2369. https://doi.org/10.1126/science.291.5512.2364

